# Hidden in plain sight - highly abundant and diverse planktonic freshwater *Chloroflexi*

**DOI:** 10.1101/366732

**Authors:** Maliheh Mehrshad, Michaela M. Salcher, Yusuke Okazaki, Shin-ichi Nakano, Karel Šimek, Adrian-Stefan Andrei, Rohit Ghai

## Abstract

**Background:** Representatives of the phylum *Chloroflexi*, though reportedly highly abundant (up to 30% of total prokaryotes) in the extensive deep water habitats of both marine (SAR202) and freshwater (CL500-11), remain uncultivated and uncharacterized. There are few metagenomic studies on marine *Chloroflexi* representatives, while the pelagic freshwater *Chloroflexi* community is largely unknown except for a single metagenome-assembled genome of CL500-11.

**Results:** Here we provide the first extensive examination of the community composition of this cosmopolitan phylum in a range of pelagic habitats (176 datasets) and highlight the impact of salinity and depth on their phylogenomic composition. Reconstructed genomes (53 in total) provide a perspective on the phylogeny, metabolism and distribution of three novel classes and two family-level taxa within the phylum *Chloroflexi*. We unraveled a remarkable genomic diversity of pelagic freshwater *Chloroflexi* representatives that thrive not only in the hypolimnion as previously suspected, but also in the epilimnion. Our results suggest that the lake hypolimnion provides a globally stable habitat reflected in lower species diversity among hypolimnion specific CL500-11 and TK10 clusters in distantly related lakes compared to a higher species diversity of the epilimnion specific SL56 cluster. Cell volume analyses show that the CL500-11 are amongst the largest prokaryotic cells in the water column of deep lakes and with a biomass:abundance ratio of two they significantly contribute to the deep lake carbon flow. Metabolic insights indicate participation of JG30-KF-CM66 representatives in the global cobalamin production via cobinamide to cobalamin salvage pathway.

**Conclusions:** Extending phylogenomic comparisons to brackish and marine habitats suggests salinity as the major influencer of the community composition of the deep-dwelling *Chloroflexi* in marine (SAR202) and freshwater (CL500-11) habitats as both counterparts thrive in intermediate brackish salinity however, freshwater habitats harbor the most phylogenetically diverse community of pelagic *Chloroflexi* representatives that reside both in epi- and hypolimnion.

## Background

In recent years, a combination of improved cultivation techniques and the use of cultivation-free approaches has led to an increasingly detailed understanding of several groups of abundant and ubiquitous freshwater microbes e.g. *Actinobacteria* [1–3], *Betaproteobacteria* [3–6], *Alphaproteobacteria* [3, 7–9] and *Verrucomicrobia* [10]. However, there are still cases of several ubiquitous groups that have largely eluded extensive characterizations. One such important instance is the phylum *Chloroflexi*, that has been shown to be abundant (up to 26% of total prokaryotic community), but mostly in the hypolimnion of lakes. In particular, the CL500-11 lineage (class *Anaerolineae*) is a significant member in deeper waters. Originally described from Crater Lake (USA) (>300m depth) using 16S rRNA clone library and oligonucleotide probe hybridization [11, 12], these microbes have been found to constitute consistently large fractions of prokaryotic communities (up to 26%) in deep lake hypolimnia all over the world [11–16]. The only genomic insights into their lifestyle come from a single metagenomic assembled genome (MAG) from Lake Michigan (estimated completeness 90%) along with *in situ* expression patterns that revealed CL500-11 to be flagellated, aerobic, photoheterotrophic bacteria, playing a major role in demineralization of nitrogen-rich dissolved organic matter in the hypolimnion [16]. Another lineage is the CL500-9 cluster [11], that was described as a freshwater sister lineage of the marine SAR202 cluster (now class ‘*Ca*. Monstramaria’) [17] but since the original discovery, there have been no further reports of its presence in other freshwater environments. Apart from these, there are only sporadic reports (of 16S rRNA sequences) for pelagic *Chloroflexi*, with little accompanying ecological information (e.g. SL56, TK10 etc.) [14, 15, 18–20].

In this work, we attempt to provide a combined genomic perspective on the diversity and distribution of *Chloroflexi* from freshwater, brackish and marine habitats. Using publicly available metagenomic data supplemented with additional sequencing from both epilimnion and hypolimnion at multiple sites, we describe three novel class-level groups of freshwater *Chloroflexi*, along with a diverse phylogenetic assortment of genomes dispersed virtually over the entire phylum. Our results also suggest that origins of pelagic *Chloroflexi* are likely from soil and sediment habitats and that their phylogenetic diversity at large correlates inversely to salinity, with freshwater habitats harboring the most diverse phylogenetic assemblages in comparison to brackish and marine habitats.

## Results and discussion

### Abundance and diversity of the phylum *Chloroflexi* in freshwater environments

Based on 16S rRNA read abundances from 117 metagenomes from lakes, reservoirs and rivers, representatives of the phylum *Chloroflexi* comprised up to seven percent of the prokaryotic community in the epilimnion (Figure 1A, 1B), however, with large fluctuations. Similar to previous observations [11–16], the CL500-11 lineage dominated hypolimnion samples (reaching at least 16% in all but one sample, and nearly 27% in one sample from Lake Biwa) (Figure 1C), apart from a lesser-known group referred to as the TK10 cluster. The majority of TK10 related 16S rRNA sequences in the SILVA database [21] originate from soil, human skin or unknown metagenomic samples, while only four (1.5%) are from freshwaters (Supplementary Figure S1A).

**Figure 1.**
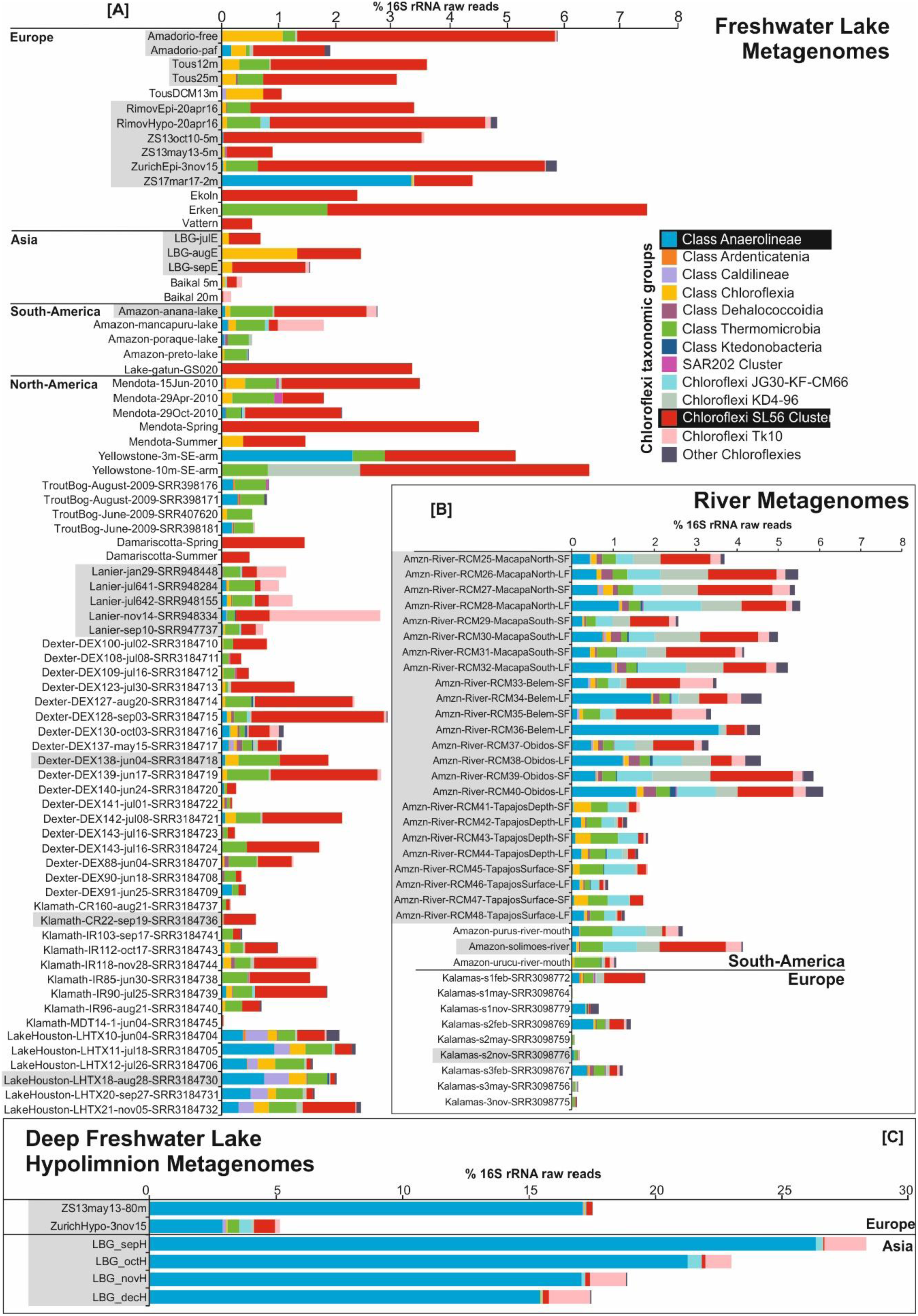
Distribution of *Chloroflexi* related 16S rRNA reads in unassembled metagenomic datasets of freshwater environments. *Chloroflexi* related 16S rRNA reads were further assigned to lower taxonomic levels based on the best BLAST to class-level taxa. Values are shown as a percentage of total prokaryotic community in [A] freshwater lakes, [B] rivers, and [C] deep lake hypolimnion. Datasets highlighted in gray were used for assembly. The complete list of datasets used and their metadata is available in Supplementary Table S3.

Surprisingly, the epilimnion samples were dominated by “SL56 marine group” (up to ca. 5% of total prokaryotic community). SL56 related sequences of SILVA have been recovered from a freshwater lake [22] and the Global Ocean Series datasets (GOS) [23]. However, the GOS sample from which they were described is actually a freshwater dataset, Lake Gatun (Panama). It is quite evident from our results (Figure 1, Supplementary Figure S2) that this cluster is consistently found only in lakes, reservoirs and rivers but not in the marine habitat, suggesting it has been incorrectly referred to as a “marine group”. Another group of sequences, referred to as JG30-KF-CM66, described from diverse environments (uranium mining waste pile, soil, freshwater, marine water column and sediment) was found to be preferentially distributed in rivers (particularly the River Amazon) than lakes (Figure 1A and B), albeit at very low abundances (maximum 1% of total prokaryotes). Similar abundances were found in the brackish Caspian Sea (depths 40m and 150m) (Supplementary Figure S2).

However, we could find no support for the presence of either the SAR202 cluster or its freshwater sister clade CL500-9 in all freshwater metagenomic datasets examined. In marine and brackish habitats, SAR202 are almost exclusively found in the dark aphotic layers, where they account for up to 30% of the prokaryotic community [24–26]. If there are any SAR202 related clades in freshwater habitats they are certainly not very abundant or perhaps did not originate from the water column in the original report [11] (Supplementary Figure S1). Overall, even though relative abundances of *Chloroflexi* in the freshwater epilimnia are far lower than in the deeper waters, they are home to a rich and widespread collection of novel groups.

With these observations, it is also readily apparent that in the aquatic environments examined here (freshwater, brackish and marine), the diversity of *Chloroflexi* representatives is substantially different, with the freshwater environments harboring a phylogenetically more diverse assortment of groups than either the brackish or the marine. Moreover, there is clear evidence for the presence of freshwater only groups (e.g. SL56), and marine and brackish only groups (SAR202), reiterating that salinity is a barrier towards microbial habitat transitions between freshwater and marine ecosystems [27]. It is by no means an insurmountable barrier as relatively recent transitions from freshwater to marine (e.g. the freshwater ‘*Ca*. Methylopumilus spp.’ and marine OM43 [28, 29]) and in reverse (marine *Pelagibacter* and freshwater LD12 [30, 31]) have both been proposed. However, it is likely that the groups found in brackish environments may perhaps be simply better “primed” for more successful forays. We do find examples of groups that are present in freshwater and brackish metagenomes (JG30-KF-CM66 and CL-500-11).

### The major freshwater *Chloroflexi* representatives

Automated binning of *Chloroflexi* related contigs from assemblies of each 57 datasets belonging to 14 different environments (26 lakes/reservoirs, 26 rivers and 3 brackish datasets) resulted in segregation of 102 MAGs (metagenome-assembled genomes) in total (Supplementary Table S1). Phylogenetic analysis of MAGs with 30% or higher completeness (n=53) shows that a remarkably high diversity of MAGs was recovered from practically all well-known *Chloroflexi* classes (Figure 2). 35 MAGs constituted three separate novel class level lineages with no available cultured representatives (SL56, TK10 and JG30-KF-CM66).

**Figure 2.**
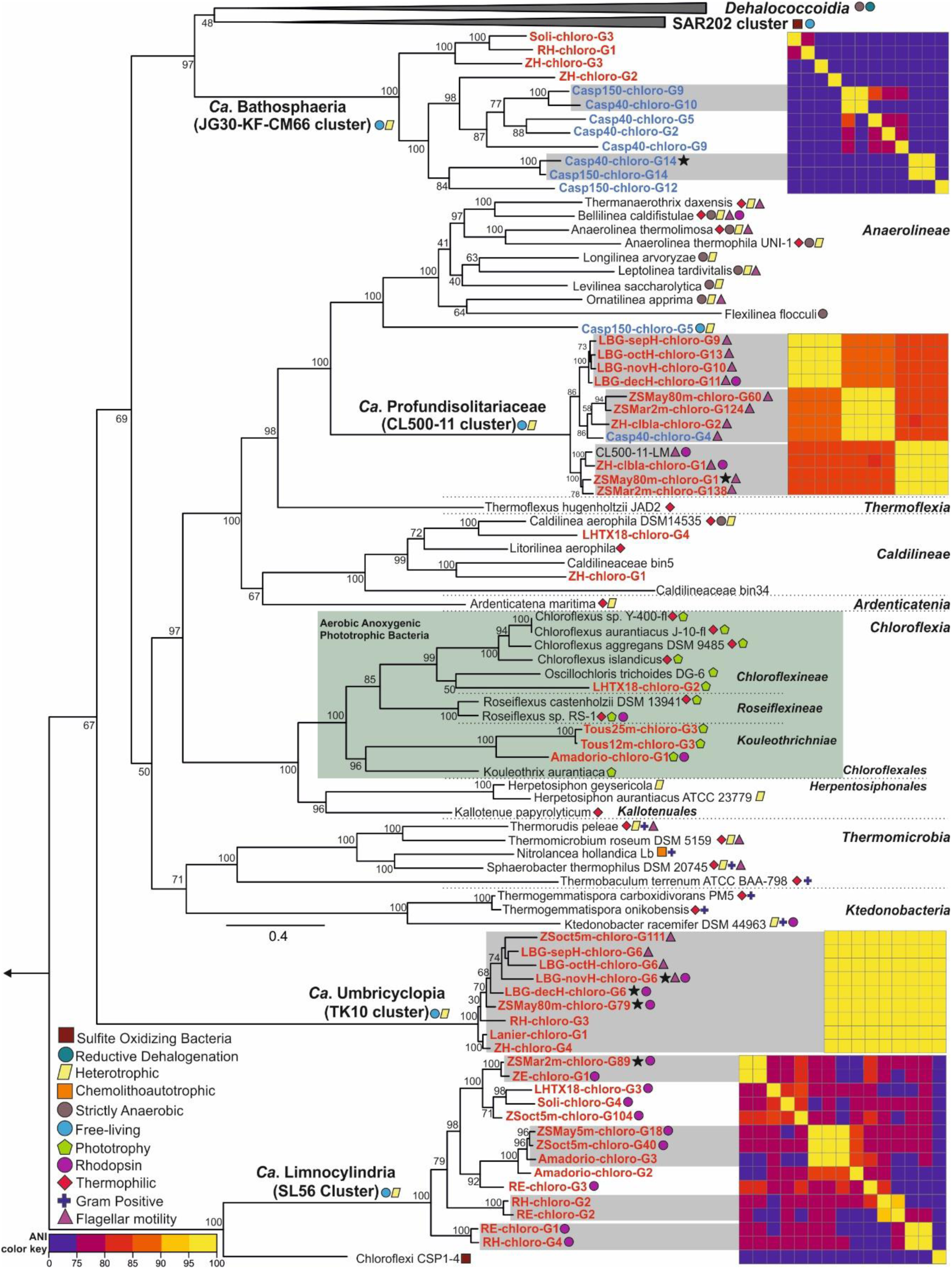
Phylogeny of the *Chloroflexi* reconstructed MAGs. Maximum likelihood phylogenomic tree reconstructed by adding the complete genomes and available MAGs of representatives from all known *Chloroflexi* classes and reconstructed MAGs of this study with the completeness higher than 30% (shown in red for freshwater originated MAGs and blue for the Caspian Sea MAGs) to the built-in tree of life in PhyloPhlAN. An asterisk next to a MAG shows the presence of 16S rRNA. Bootstrap values (%) are indicated at the base of each node. Legends for lifestyle hints are on bottom left. Average nucleotide identity comparison (ANI) heat map for MAGs of each cluster is shown to the right of each cluster. Reconstructed genomes belonging to the same species are shown inside a grey box. A color key for the ANI is shown at the bottom left. The green box shows the Aerobic anoxygenic phototrophic members of the class *Chloroflexia*.

While CARD-FISH detected high numbers of the CL-500-11 cells in Lake Zurich epilimnion during partial mixis in winter, peak abundance levels were always found in deeper zones, in both Lake Zurich (up to 11% of all prokaryotes; Figure 3A) and Lake Biwa (up to 14%; Figure 3D). CL500-11 abundance correlated negatively with both temperature and chlorophyll *a* concentration (Supplementary Figure S3). In the Řimov reservoir samples however, CL-500-11 was below the detection limit (<0.18%), suggesting that this relatively shallow habitat (maximum depth 43m) does not represent a preferred niche for this group of bacteria (Supplementary Figure S4). CL-500-11 cells have been previously visualized by CARD-FISH and shown to be large, curved cells [13]. Similar shapes and sizes were observed in FISH samples from Lake Zurich with mean lengths of 0.92 μm (range 0.4-1.6 μm; n=277) and widths of 0.28 μm (range 0.19-0.39 μm). Analyzing the cell volumes (0.06 μm^3^ median) and biomass for this cluster in comparison to all prokaryotes (Figure 3C) suggests an extremely high contribution of the CL-500-11 population to total microbial biomass. Their biomass:abundance ratio is nearly 2, i.e. at 10% abundance they comprise almost 20% of the total prokaryotic biomass, indicating a remarkable adaptation to the relatively oligotrophic deep hypolimnion, attaining high populations even with their large cell sizes.

**Figure 3.**
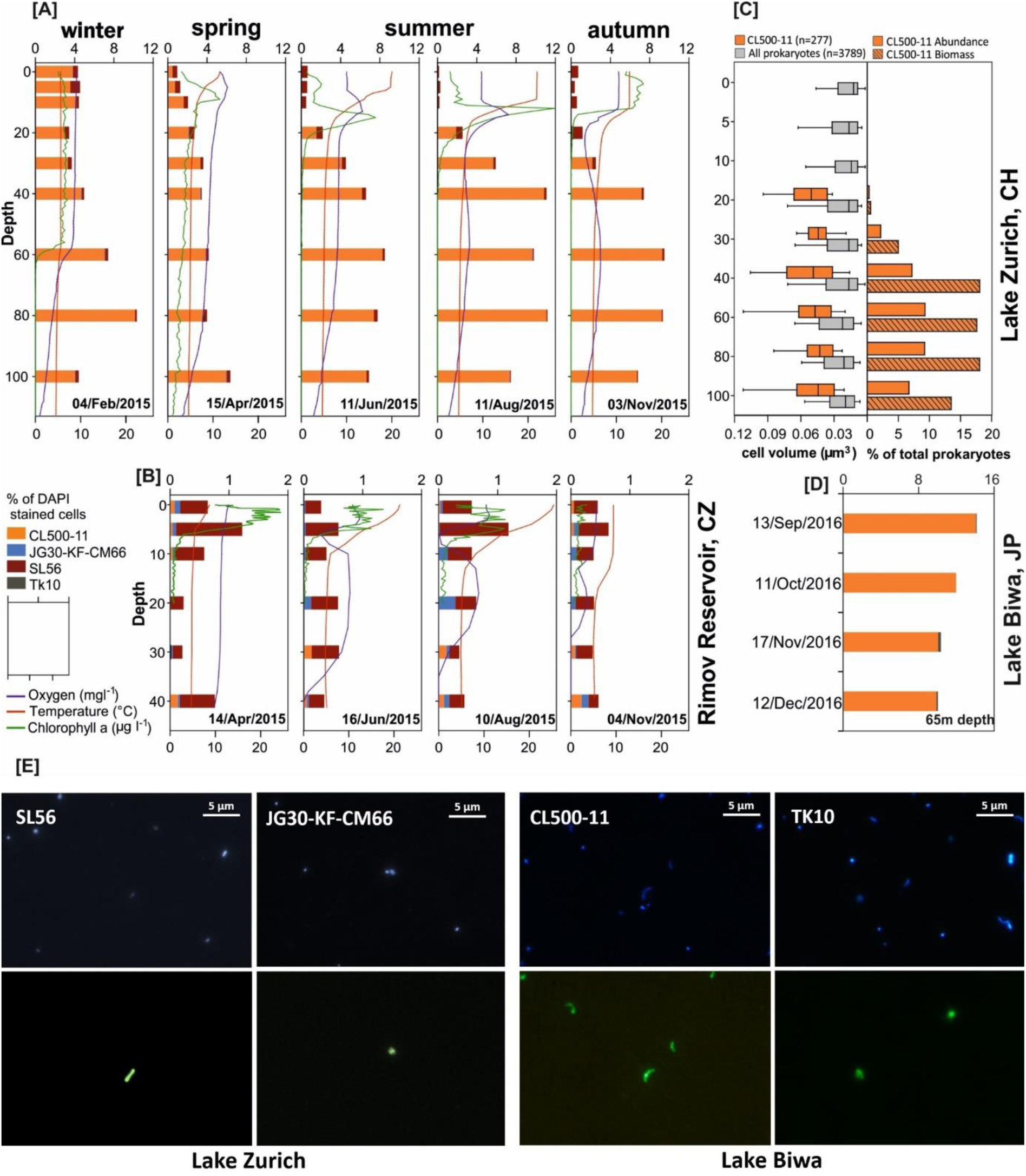
Spatiotemporal distribution and cell shape of different *Chloroflexi* lineages based on CARD-FISH analysis. Seasonal dynamic and vertical stratification of different *Chloroflexi* lineages according to CARD-FISH analysis in [A] Lake Zurich at five sampling times and [B] Rimov reservoir at four sampling times during the year 2015. The stacked bars show the percentage of DAPI stained cells (top axis) and the smooth lines show vertical profiles of water temperature, oxygen and chlorophyll a (bottom axis). [C] Cell volume (μm3) of CARD-FISH stained Chloroflexi CL500-11 (n=277) and all prokaryotes (n=3789) along depth profile of the Lake Zurich on November 3rd 2015. Boxes show 5th and 95th percentile and the vertical line represents the median. The percentage of CL500-11 abundance and biomass among prokaryotes of the same depth profile is shown on the right side. [D] The abundance of Chloroflexi lineages in 65m depth of the Lake Biwa at four sampling times in 2016. [E] CARD-FISH images of different *Chloroflexi* lineages. An identical microscopic field is shown for each column, with the DAPI-stained cells in the top and bacteria stained by cluster specific CARD-FISH probes of each cluster on the bottom. The scale is shown on the top right side of the DAPI stained cells field.

We recovered 11 MAGs (10 freshwaters, 1 brackish) for CL500-11 in total. All four MAGs of Lake Biwa from different months form a single species. However, the two species from Lake Zurich appear to coexist throughout the year (March, May and November) with one species branching together with the previously described MAG from Lake Michigan (CL500-11-LM) [16], and the other species having close representatives also in the brackish Caspian (>95% ANI) and similar metagenomic fragment recruitment patterns (Figure 2 an 4C). We propose the candidate genus Profundisolitarius (Pro.fun.di.so.li.ta’ri.us. L. adj. profundus deep; L. adj. solitarius alone; N.L. masc. n. Profundisolitarius a sole recluse from the deep) within *Candidatus* Profundisolitariaceae fam. nov. for the CL500-11 cluster (class *Anaerolinea*).

On the other hand, the SL56 group is the dominant lineage in the Řimov reservoir (maximum 1.1%), both by 16S rRNA and CARD-FISH analyses (Figure 1 and Figure 3). Maximal abundances were nearly always found at around 5-20m at temperatures of ca. 15ºC, suggesting that this group is primarily epilimnetic (Supplementary Figures S3 and S4). This region of the water column (thermocline), apart from having a temperature gradient, also has significantly lower light intensity in comparison to surface layers. Peak abundances of the low light adapted cyanobacterium *Planktothrix rubescens* [32] at around 13m depth in the stratified summer profiles of Lake Zurich, coincide with maximal abundances of the SL56 (Supplementary Figure S3). SL56 cells are rod-shaped and elongated (average length=0.68 ± 0.25 μm; average width=0.35 ± 0.09 μm; n=6; Figure 3E). To the best of our knowledge, this is the first report of a freshwater specific *Chloroflexi* group that appears to thrive in the epilimnion.

A total of 14 MAGs were recovered for SL56 cluster (1 containing 16S rRNA) and form a class level lineage, considerably divergent from all known *Chloroflexi* (Figure 2). Their sole relative is a single MAG (*Chloroflexi* CSP1-4) described from aquifer sediment [33]. The 16S rRNA clade to which the CSP1-4 reportedly affiliates to is Gitt-GS-136 [33] and the majority of sequences in this clade originate from either soil or river sediments (information from SILVA taxonomy). However, we were unable to detect any 16S rRNA sequence (partial or complete) in the available genome sequence of CSP1-4. The next closest clade (in the 16S rRNA taxonomy) to Gitt-GS-136 and SL56 is KD4-96, whose sequences were obtained from the same habitats (See Supplementary Figure S1B). In addition, all known 16S rRNA sequences from the SL56 group originate only from freshwaters (Lake Gatun, Lake Zurich etc.). Taken together, it appears that the closest phylogenetic relatives of the freshwater SL56 lineage inhabit soil or sediment habitats.

SL56 MAGs were reconstructed from geographically distant locations (Europe, North and South America, Figure 2) and at least nine different species could be detected (ANI, Figure 1). No MAGs were obtained from Lake Biwa samples but three 16S rRNA sequence were retrieved in unbinned contigs. The reconstructed MAGs are globally distributed along the freshwater datasets from the epilimnion (none detected in the deep hypolimnion) (Figure 4 and Supplementary Figure S6). No SL56 MAGs were reconstructed from the Caspian Sea and none of the recovered genomes recruited from brackish metagenomes. We propose the candidate genus Limnocylindrus (Lim.no.cy.lin’drus. Gr. fem. n. limne a lake; L. masc. n. cylindrus a cylinder; N.L. masc. n. Limnocylindrus a cylinder from a lake) within Limnocylindraceae fam. nov., Limnocylindrales ord. nov., and Limnocylindria classis. nov. for the *Chloroflexi* SL56 cluster.

**Figure 4.**
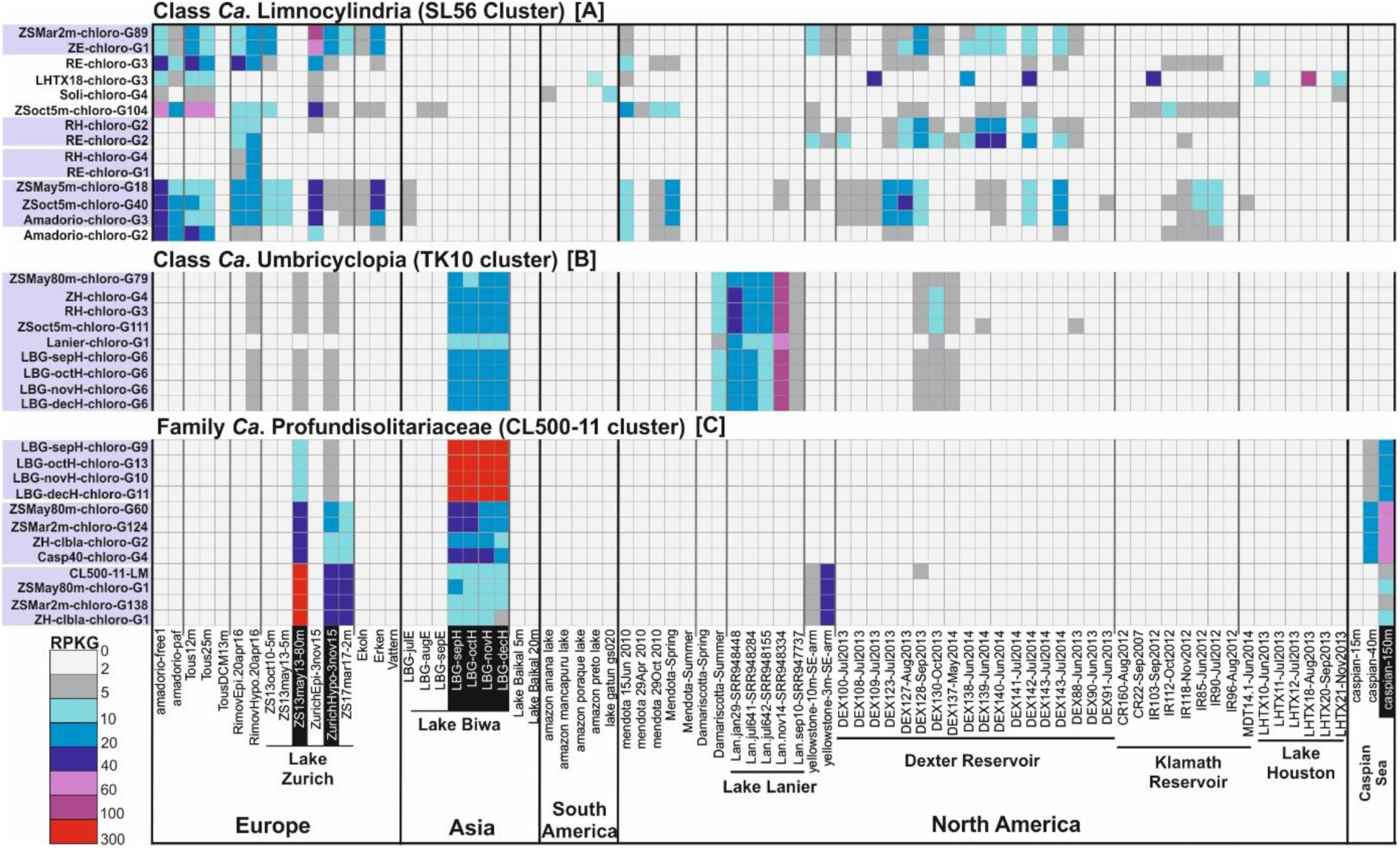
Distribution of *Chloroflexi* reconstructed MAGs in freshwater and brackish environments. The recruitment (RPKG) distribution of reconstructed MAGs of *Chloroflexi* cluster SL56 [A], TK10 [B], and CL500-11 [C] against freshwater and brackish datasets. Freshwater datasets belong to the lakes and reservoirs from Europe (16), Asia (9), South (5) and North America (47) and brackish datasets include three depths (15m, 40m, and 150m) datasets of the Caspian Sea (complete list of datasets used and their metadata is available in Supplementary Table S3). The hypolimnion datasets of Lake Zurich, Lake Biwa, and Caspian Sea are shown in black boxes. Genomes belonging to the same species are shown in a gray box.

TK10 16S rRNA sequences were found at highest abundances in Lake Biwa hypolimnion samples (maximum ca. 2%) (Figure 1A and C). Cells were ovoid with an estimated length of 1.08 ± 0.1 μm and width of 0.84 ± 0.09 μm (n=12; Figure 3E). A coherent cluster of nine MAGs (3 containing 16S rRNA Supplementary Figure S1) from geographically distant locations (Europe, Asia and North America) was recovered. These remarkably cosmopolitan organisms thriving in deeper lake strata are not very diverse (ANI values >95%). This apparent low diversity might be a consequence of a very specialized niche or what is more likely, an outcome of a relatively recent transition to freshwater, similar to ‘*Ca*. Fonsibacter’ (LD12 *Alphaproteobacteria*) [8]. No 16S rRNA representatives were detected confidently in marine or brackish metagenomes though some 16S rRNA sequences of SILVA database have been obtained from marine sediments and water column (Supplementary Figure S1). Closest relatives from 16S rRNA appear to be either from soil or sediment samples suggesting that these might be their original habitat. Interestingly, the TK10 cluster is also deep branching, only after SL56 and CSP1-4 in the phylogenetic tree of *Chloroflexi* at large, and all other *Chloroflexi* representatives (MAGs or isolate genomes) appear to be descended from a branch distinct to both of these. We suggest the candidate genus Umbricyclops (Um.bri.cy’clops. L. fem. N. umbra shadow; L. masc. n. cyclops (from Gr. Round eye; Cyclops) a cyclops; N.L. masc. n. Umbricyclops a round-eye living in the shade) within Umbricyclopaceae fam. Nov., Umbricyclopales ord. nov., and Umbricyclopia classis. nov. for this group of organisms.

CARD-FISH results show that JG30-KF-CM66 cells are spherical with an estimated diameter of 0.56 μm (± 0.15 μm; n=8; Figure 3E) however, very low proportions (<0.28%) were observed for JG30-KF-CM66 in Lake Zurich and the Řimov Reservoir depth profiles (Supplementary Figures S3 and S4). We obtained 12 MAGs, mostly from deep water column (8 brackish, 4 freshwater), one with a near complete 16S sequence, that formed a novel class level lineage in the phylogenomic analysis (Figure 1). The closest relatives of these MAGs are marine SAR202 and *Dehalococcoidea* (Figure 1 and Supplementary Figure S1). Within this cluster distinct groups of brackish and freshwater MAGs can be distinguished. We suggest the candidate genus Bathosphaera (Ba.tho.sphae’ra. Gr. adj. bathos deep; L. fem. n. sphaera a sphere; N.L. fem. n. Bathosphaera a coccoid bacteria living in the deep) within Bathosphaeraceae fam. nov., Bathosphaerales ord. nov., and Bathosphaeria classis. nov. for the *Chloroflexi* JG30-KF-CM66 cluster.

We also recovered MAGs in the classes *Chloroflexia* (4 MAGs) and *Caldilineae* (2 MAGs) (Figure 1). *Chloroflexia* MAGs were related to mesophilic *Oscillochloris trichoides* DG-6 in sub-order *Chloroflexineae* (1 MAG) and 3 other MAGs to *Kouleothrix aurantiaca* in the *Kouleotrichaceae* fam. nov. forming a new sub-order for which we propose the name *Kouleothrichniae* sub-order. nov. None of these MAGs show any significant fragment recruitment apart from their place of origin. An additional 14 MAGs from the Caspian affiliated to the SAR202 cluster which will not be further discussed here as they have already been described [26].

### Contribution of freshwater *Chloroflexi* in ecosystem functioning

Metabolic insights into the reconstructed *Chloroflexi* MAGs (completeness ≥30%) suggest a primarily heterotrophic life style which in some groups is boosted by light driven energy generation either via rhodopsins (CL500-11, *Chloroflexales*, SL56, and TK10) or aerobic anoxygenic phototrophy (*Chloroflexales*). The MAGs of each cluster contain necessary genes for central carbohydrate metabolism including glycolysis, gluconeogenesis, and tricarboxylic acid cycle. Key genes for assimilatory sulfate reduction (3′-phosphoadenosine 5′-phosphosulfate (PAPS) synthase and sulfate adenylyltransferase) were absent in most MAGs suggesting the utilization of exogenous reduced sulfur compounds [34]. Denitrification genes (nitrate reductase/nitrite oxidoreductase alpha and beta subunits and nitrite reductase) were found in TK10 MAGs but the subsequent enzymes responsible for the production of molecular nitrogen were absent.

In aquatic environments *Thaumarchaeota* and *Cyanobacteria* are the main source of cobalamin and its corrinoid precursors for the large community of auxotrophs or those few capable of salvage [35, 36]. De-novo synthesis of cobalamin has a high metabolic cost, and the Black Queen Hypothesis has been put forward as an explanation for reasons why only a few community members undertake its production [35, 37, 38]. None of the reconstructed *Chloroflexi* MAGs encode necessary genes for corrin ring biosynthesis from scratch and high affinity cobalamin (BtuBFCD) or other suspected corrinoid (DET1174-DET1176) [39] transporters were also missing which may be a consequence of genome incompleteness or use of an undescribed transporter. However, not all these organisms seem to be auxotrophs as the MAGs of JG30-KF-CM66 cluster encode genes for cobinamide to cobalamin salvage pathway that utilizes imported corrinoids together with intermediates from the riboflavin biosynthesis pathway to synthesize cobalamin [40]. ZH-chloro-G3 MAG contains an almost complete cobalamin salvage (only missing CobC) and riboflavin biosynthesis pathway (Supplementary Table S2).

Flagellar assembly genes were present in several MAGs of CL500-11 and TK10 clusters (Figure 1 and Supplementary Table S2). However, the L and P-ring components that anchor flagella to the outer membrane were missing in all flagellated MAGs and reference *Chloroflexi* genomes (e.g. *Thermomicrobium* [41], *Sphaerobacter* [42]). In addition, MAGs and reference *Chloroflexi* genomes did not encode genes for LPS biosynthesis and no secretion systems, apart from Sec and Tat were detected (Type I – IV secretion systems that are anchored in the outer membrane are absent) (Supplementary Table S2). Taken together the comparative genomics of available *Chloroflexi* genomes bolster inferences that while electron micrographs suggest two electron dense layers in most members of this phylum, *Chloroflexi* likely possess a single lipid membrane (monoderm) rather than two (diderms) [42].

Rhodopsin-like sequences were recognized in 18 MAGs of this study from representatives of CL500-11, *Chloroflexia*, SL56, and TK10 that are phylogenetically closest to xanthorhodopsins (Supplementary Figure S8A and B), and are tuned to absorb green-light similar to other freshwater and coastal rhodopsins [2, 23] (Supplementary Figure S8C). Several MAGs encode genes for carotenoid biosynthesis allowing the possibility of a carotenoid antenna that is the hallmark of xanthorhodopsins [43–45]. Of the residues involved with binding salinixanthin (the predominant carotenoid of *Salinibacter ruber*), we found a surprisingly high number conserved (10 identical out of 12 in at least one rhodopsin sequence) (Supplementary Figure S8D), suggesting that a carotenoid antenna may be bound, making at least some of these sequences *bonafide* xanthorhodopsins.

Even representatives of CL500-11 and TK10 that are primarily found in the hypolimnion during stratification are capable of phototrophy, however, they can potentially access the photic zone during winter and early spring mixis. Apart from rhodopsin-based photoheterotrophy, we also retrieved MAGs of the class *Chloroflexia* encoding genes for photosystem type II reaction center proteins L and M (pufL and pufM), bacteriochlorophyll and carotenoid biosynthesis. The pufM gene sequences cluster together with other *Chloroflexi*-related pufM sequences (Supplementary Figure S9). However, no evidence for carbon fixation, either via the 3-hydroxypropionate pathway or the Calvin-Benson cycle was found in any photosystem bearing MAG which might be a consequence of MAG incompleteness. It may also be that these are aerobic anoxygenic phototrophs that do not fix carbon e.g. freshwater *Gemmatimonadetes* and *Acidobacteria* (both aerobic) [46].

### Evolutionary history of pelagic Chloroflexi

It is apparent from the phylogenomic analyses that the collection of representatives of the phylum *Chloroflexi* recovered in this work, along with the existing genome sequences from isolates and MAGs, offers only a partial sketch of the complex evolutionary history of the phylum at large. For example, the most divergent branches ‘*Ca*. Limnocylindria’ (SL56 cluster) and ‘*Ca*. Umbricyclopia’ (TK10 cluster) have practically no close kin apart from an aquifer sediment MAG (related to ‘*Ca*. Limnocylindria’). However, related 16S rRNA clones have been recovered from soil/sediments for both these groups, suggesting transitions to a pelagic lifestyle. Factoring the absence of related marine 16S rRNA sequences for these groups, in addition to their undetectability in marine metagenomic datasets also suggests an ancestry from soil/sediment rather than the saline environment. While the possibility of a marine origin cannot be formally excluded, the directionality of a transition from soil/sediment to freshwater water columns appears most likely. Moreover, given that ‘*Ca*. Limnocylindria’ and ‘*Ca*. Umbricyclopia’ diverge prior to the divergence of the classes *Dehalococcoidea* and marine SAR202 (class ‘*Ca*. Monstramaria’), which are the only ecologically relevant marine *Chloroflexi* known as yet (the former in marine sediments and the latter in deep ocean water column), it is likely that ancestral *Chloroflexi* originated in a soil/sediment habitat. The success of marine SAR202 in the deep oceans is remarkable, it is the most widely distributed, perhaps numerically most abundant *Chloroflexi* group on the planet. However, some 16S rRNA sequences from its closest relatives, *Dehalococcoidea*, have also been recovered from freshwater sediments, even though the vast majority appear to be from deep marine sediments (both anoxic habitats).

In this study, we significantly expand our conceptions regarding the diversity of pelagic *Chloroflexi* and their possible origins from soil/sediment habitats. Similar evolutionary trajectories are beginning to be visible for other freshwater microbes, e.g. the closest relatives of freshwater *Actinobacteria* (‘*Ca*. Nanopelagicales’ [2]) being soil *Actinobacteria* or the transition of methylotrophic *Betaproteobacteria* (‘*Ca*. Methylopumilus’) from sediments to the water column [4, 47], and as more and more prokaryotic groups are examined and the study is expanded to the sediment and soil habitats we will finally be able to reconstruct the sequence of events that have led to the complex mosaic of freshwater microbial communities as we see them today.

## Methods

### Sample collection

#### Řimov reservoir

Representative water samples of epilimnion (0.5m) and hypolimnion (30m) were taken on April 20^th^ 2016 from this mesoeutrophic reservoir (South Bohemia, Czech Republic). The sampling site is located at the deepest part (43m) of the reservoir 250m from the dam. For more detail about the reservoir see the reference [48].

#### Lake Zurich

Samples from this oligo-mesotrophic Lake (Switzerland) were collected on October 13^th^ 2010 (5m depth), May 13^th^ 2013 (5m and 80m depth), November 3^rd^ 2015 (5m and 40-80m depth), and March 17^th^ 2017 (2m depth). The sampling site is located at the deepest part (136m) of Lake Zurich.

#### Lake Biwa

Samples from this mesotrophic Lake were collected at a pelagic station (35° 12’58” N 135° 59’55” E; water depth = ca. 73m) in 2016. Samples from the epilimnion (5m depth) were taken on July 20^th^, August 18^th^, and September 27^th^. Samples from the hypolimnion (65m) were taken on September 13^th^, October 11^th^, November 17^th^, and December 12^th^.

All water samples were sequentially pre-filtered through 20 and 5 μm pore-size filters and the flow-through microbial community was concentrated on 0.22 μm filters (polycarbonate (PCTE) membrane filters, Sterlitech, USA, for Řimov and Zurich samples and polyethersulfone filter cartridges (Millipore Sterivex SVGP01050) for Lake Biwa samples. DNA extraction of Řimov reservoir and Lake Zurich samples was performed using the standard phenol-chloroform protocol [49]. For samples from Lake Biwa, DNA was extracted by PowerSoil DNA Isolation Kit (MoBio Laboratories, Carlsbad, CA, USA). Sequencing of the samples from the Řimov reservoir (n=2) and Lake Zurich (n=2) was performed using Illumina HiSeq4000 (2×151bp, BGI Genomics, Hong Kong, China), additional samples from Lake Zurich (n=4) were sequenced using Illumina HiSeq2000 (2×150X bp, Functional Genomics Center, Zurich, Switzerland) and Lake Biwa samples (n=7) were sequenced using MiSeq (2×300bp, Bioengineering Lab. Co., Ltd. Kanagawa, Japan).

Basic metadata (sampling date, latitude, longitude, depth, bioproject identifiers, SRA accessions), and sequence statistics (number of reads, read length, dataset size) of all metagenomes generated in this study are provided in Supplementary Table S3.

### Unassembled 16S rRNA read classification

A non-redundant version of the SILVA_128_SSURef_NR99 database [21] was created by clustering its 645’151 16S rRNA gene sequences into 7’552 sequences at 85% nucleotide identity level using UCLUST [50]. Ten million reads from each dataset were compared to this reduced set and an e-value cutoff of 1e-5 was used to identify candidate 16S rRNA gene sequences. If a dataset had less than 10 million reads, all reads from the dataset were used to identify candidate sequences. These candidate sequences were further examined using ssu-align, and segregated into archaeal, bacterial, and eukaryotic 16S/18S rRNA or non-16S rRNA gene sequences [51]. The bona fide prokaryotic 16S rRNA sequences were compared to the complete SILVA database using BLASTN [52] and classified into a high level taxon if the sequence identity was ≥80% and the alignment length was ≥90 bp. Sequences failing these thresholds were discarded. The 16S rRNA reads belonging to the phylum *Chloroflexi* were furtherly segregated to lower taxonomic levels of the SILVA taxonomy.

### Assembled 16S rRNA sequences from the freshwater metagenomes and 16S rRNA gene phylogeny

Assembled 16S rRNA sequences of the 120 assembled freshwater datasets were identified using Barrnap with default parameters (https://github.com/tseemann/barrnap). Genes encoding 16S rRNA were aligned using the SINA web aligner [53], imported to ARB [54] using the SILVA_128_SSURef_NR99 database [21], manually checked, and bootstrapped maximum likelihood trees (GTR-GAMMA model, 100 bootstraps) were calculated with RAxML [55].

### Collection of depth profile samples for CARD-FISH analyses

Řimov Reservoir was sampled four times in 2015, during the spring phytoplankton bloom (April 14^th^), early summer (June 16^th^), late summer (August 10^th^), and autumn (November 04^th^). Vertical profiles of physicochemical parameters were taken by a YSI multiprobe (Yellow Springs Instruments, model 6600, Yellow Springs, OH, USA) and profiles of different phytoplankton groups differentiated by their fluorescent spectra were obtained with a fluorescence probe (FluoroProbe, TS-16-12, bbe Moldaenke GmbH, Schwentinental, Germany). Water samples were taken from 0, 5, 10, 20, 30, and 40m depths (n=28).

Lake Zurich was sampled five times in 2015, during winter mixis (February 4^th^), the spring phytoplankton bloom (April 15^th^), early summer (June 11^th^), late summer (August 11^th^), and autumn (November 03^th^). Sampling included vertical profiles of physicochemical parameters using a YSI multiprobe (Yellow Springs Instruments, model 6600, Yellow Springs, OH, USA) and profiles of four phytoplankton groups (*Planktothrix rubescens*, green algae, diatoms and cryptophytes) differentiated by different fluorescent spectra using a submersible fluorescence probe (FluoroProbe, TS-16-12, bbe Moldaenke GmbH, Schwentinental, Germany). Water samples for bacterial analyses were taken from 0, 5, 10, 20, 30, 40, 60, 80, and 100m (n=45).

CARD-FISH samples from Lake Biwa were taken at the same occasion as the metagenomic samples. In the present study, only the hypolimnetic samples were analyzed (September, October, November, and December 2016 at 65 m depth)

### Design and application of novel specific 16S rRNA probes for different *Chloroflexi* clusters

CARD-FISH (fluorescence *in situ* hybridization followed by catalyzed reporter deposition) with fluorescein-labeled tyramides was conducted as previously described [56] with a probe specific for the CL500-11 cluster of *Chloroflexi* [13] and three novel probes targeting the lineages SL56, JG30-KF-CM66, and TK10 (see Supplementary Table S4 for details). A total of 54 16S rRNA sequences from multiple groups of freshwater *Chloroflexi* (e.g. CL500-11, SL56, TK10, and JG30-KF-CM66, Supplementary Figure S1A), were extracted from MAGs (n=7) or unbinned *Chloroflexi* contigs (n=47). These additional sequences were used to supplement a local reference database for prokaryotes (see methods) and design FISH probes for these groups. Probe design based on 16S rRNA genes was done in ARB [54]. A bootstrapped maximum likelihood tree (GTR-GAMMA model) of 16S rDNA sequences (Supplementary Figure S1) served as backbone for probe design with the ARB tools probe_design and probe_check. The resulting probes with their corresponding competitor and helper oligonucleotides (Supplementary Table S4) were tested with different formamide concentrations to achieve stringent hybridization conditions. CARD-FISH stained samples were analyzed by fully automated high-throughput microscopy [56]. Images were analyzed with the freely available image analysis software ACMEtool 216 (technobiology.ch), and interfering autofluorescent cyanobacteria or debris particle were individually excluded from hybridized cells. At least 10 high quality images or >1000 DAPI stained bacteria were analyzed per sample. Cell sizes of CARD-FISH stained *Chloroflexi* CL500-11 and all prokaryotes were measured from one depth profile from Lake Zurich (November 3^rd^ 2015) with the software LUCIA (Laboratory Imaging Prague, Czech Republic) following a previously described workflow [57]. At least 200 individual DAPI stained cells (corresponding to 24-65 CL500-11 cells) per sample were subjected to image analysis. Total numbers of heterotrophic prokaryotes were determined by an inFlux V-GS 225 cell sorter (Becton Dickinson) equipped with a UV (355nm) laser. Subsamples of 1 ml were stained with 4’,6-Diamidino-2-phenylindole (DAPI, 1 μg ml^-1^ final concentration), and scatter plots of DAPI fluorescence vs. 90° light scatter were analyzed with an in-house software (J. Villiger, unpublished).

### Metagenome assembly

Lake Biwa (7 datasets) and Lake Zurich (4 datasets) were assembled using metaSPAdes (-k 21,33,55,77,99,127)[58]. All other datasets, including those from the Řimov Reservoir, were assembled using megahit (--k-min 39 --k-max 99/ 151 --k-step 10 --min-count 2). A complete list of all metagenomic datasets assembled in this study (n=57) is shown in Supplementary Table S1. Prior to assembly, all datasets were quality trimmed either using sickle (https://github.com/najoshi/sickle, default parameters), or for Lake Zurich and Lake Biwa metagenomes, Trimmomatic [59] was used to remove adaptor sequences, followed by 3’ end quality-trim using PRINSEQ [60] (quality threshold = 20; sliding window size = 6) (also indicated in the Supplementary Table S3).

### Gene prediction and taxonomic analyses

Prodigal (in metagenomic mode) was used for predicting protein coding genes in the assembled contigs [61]. All predicted proteins were compared to the NCBI-NR database using MMSeqs2 (e-value 1e-3) [62] to ascertain taxonomic origins of assembled contigs.

### Metagenomic assembled genome (MAG) reconstruction

Only contigs longer than 5 kb were used for genome reconstructions. A contig was considered to belong to the phylum *Chloroflexi* if a majority of its genes gave best hits to this phylum. Chloroflexi affiliated contigs within each dataset were grouped based on the tetra-nucleotide frequencies and contig coverage pattern in different metagenomes using MetaBAT with “superspecific” setting [63]. Preliminary genome annotation for all bins was performed using Prokka [64]. Additional functional gene annotation for all *Chloroflexi* bins was performed by comparisons against COG hmms [65] using and e-value cutoff of 1e-5, and TIGRfams models [66] (using trusted score cutoffs --cut_tc) using the hmmer package [67]. The assembled genomes were also annotated using the RAST server [68] and BlastKOALA [69]. Enzyme EC numbers were predicted using PRIAM [70].

### Genome quality check, size estimation and phylogenomics

CheckM [71] was used to estimate genome completeness. A reference phylogenomic tree was made by inserting complete genomes of representatives from all known *Chloroflexi* classes and reconstructed MAGs of this study (with estimated completeness of 30% and higher) to the built-in tree of life in PhyloPhlAN [72]. PhyloPhlAN uses USEARCH [50] to identify the conserved proteins and subsequent alignments against the built-in database are performed using MUSCLE [73]. Finally, an approximate maximum-likelihood tree is generated using FastTree [74] with local support values using Shimodaira–Hasegawa test [75]. This analysis confirmed that all reconstructed MAGs belong to the phylum *Chloroflexi* and also suggests their phylogenetic affiliations within the phylum.

### Metagenomic fragment recruitment

To avoid bias in abundance estimations owing to the presence of highly related rRNA sequences in the genomes/metagenomes, rRNA sequences in all genomes were masked. After masking, recruitments were performed using BLASTN [52], and a hit was considered only when it was at least 50 bp long, had an identity of >95% and an e-value of ≤1e-5. These cutoffs approximate species-level divergence [76]. These hits were used to compute the RPKG (reads recruited per kilobase of genome per gigabase of metagenome) values that reflect abundances that are normalized and comparable across genomes and metagenomes of different sizes.

### Single gene phylogeny and average nucleotide identity (ANI)

The pufM and rhodopsin protein sequence alignments were performed using MUSCLE [73], and FastTree2 [74] was used for creating the maximum-likelihood tree (JTT+CAT model, gamma approximation, 100 bootstrap replicates). Average Nucleotide Identity (ANI) was calculated as defined in [76].

### Availability of supporting data The

The metagenomic Raw read files of the epilimnion and hypolimnion of Řimov reservoir, Lake Zurich and Lake Biwa are archived at the DDBJ/EMBL/GenBank and can be accessed under the Bioprojects PRJNA429141, PRJNA428721 and PRJDB6644 respectively. All assembled genomic bins of this study can be accessed under the Bioproject PRJNA356693.

### Ethics approval

Ethics approval was not required for the study.

## Author contributions

M.M. and R.G. concieved and designed the research. M.M., Y.O., S.H.N, M.M.S., R.G., K.S., were involved in sampling, sample processing and filtration. M.M., M.M.S., Y.O., A.S.A and R.G. performed metagenomic data analyses. M.M.S. and Y.O. performed CARD-FISH analyses. M.M, M.M.S. and R.G wrote the manuscript with input from all authors. All authors read and approved the final text.

## Acknowledgements

The authors thank P. Znachor, P. Rychtecký, T. Shabarova, and J. Nedoma for help with sampling of the Římov Reservoir and E. Loher and T. Posch for help with sampling of Lake Zurich. S. Neuenschwander is acknowledged for help with metagenomic library preparation of Lake Zurich. Aharon Oren is acknowledged for taxa nomenclature review. M.M. was supported by the Czech Academy of Sciences (Postdoc program PPPLZ application number L200961651). R.G., A.S.A. and K.S. and were supported by the research grants 17-04828S and 13-00243S from the Grant Agency of the Czech Republic. The collaborative work of M.M., Y.O., S.H.N., and K.S. was supported by JSPS Bilateral Joint Research Project No. JSPS-17-17. Y.O. and S.H.N. were supported by Environment Research and Technology Development Fund No 5-1607 of the Ministry of the Environment, Japan. Y.O. was also supported by JSPS KAKENHI Grant No 15J00971 and by The Kyoto University Foundation. Computation time was partially provided by the Super Computer System, Institute for Chemical Research, Kyoto University.

---

**Supplementary Figure S1:**
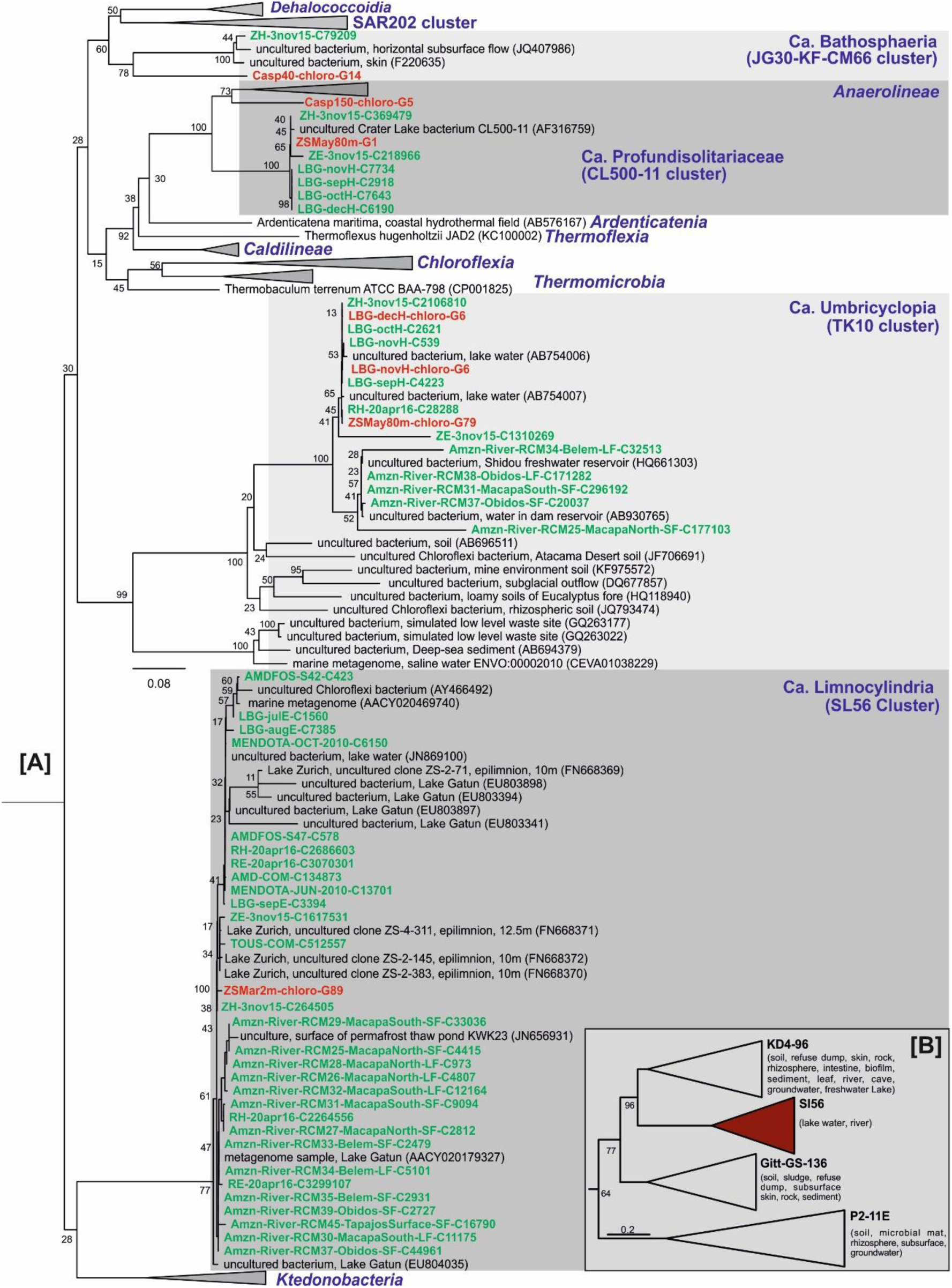
Maximum likelihood 16S rRNA tree reconstructed by adding the 16S rRNA sequences assembled from freshwater metagenomes to existing sequences of the SSURef_NR99_128 database in the phylum *Chloroflexi*. Bootstrap values (%) are indicated at the base of each node. 16S rRNA sequences present in a MAG are highlighted in red and the other metagenomic assembled 16S rRNA sequences are highlighted in green [A]. Maximum likelihood 16S rRNA tree of the SL56 cluster together with its closely related clusters. The origin of the 16S rRNA sequences present in SILVA for each cluster are summarized in parenthesis [B].

**Supplementary Figure S2:**
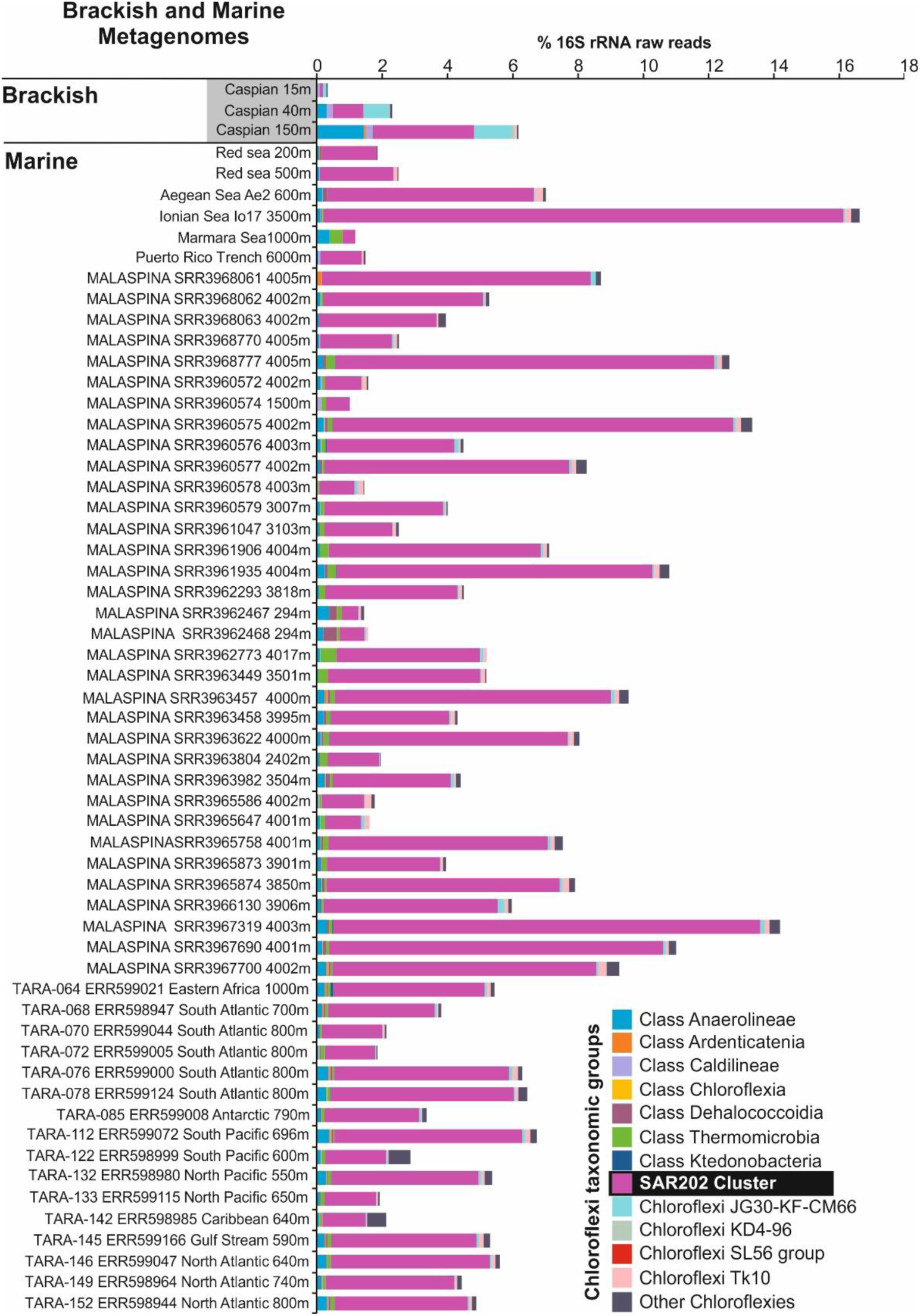
Percentage and distribution of *Chloroflexi* related 16S rRNA reads (as % of total prokaryotic community) based on unassembled metagenomic datasets in brackish and marine datasets. Brackish datasets include three different depths of the Caspian Sea. Marine datasets include Aegean Sea (one DCM and one deep dataset), Ionian Sea (one DCM and one deep dataset), Atlantic BATS, Pacific HOTS and Red Sea depth profile datasets together with selected deep datasets from MALASPINA and TARA expeditions and the Puerto Rico deep trench dataset. Chloroflexi related reads were further assigned to lower taxonomic levels of the phylum Chloroflexi based on the best BLAST hit to class-level taxa. The complete list of datasets used is available in (Mehrshad *et al*., 2017). Datasets highlighted in gray were used for the assembly.

**Supplementary Figure S3:**
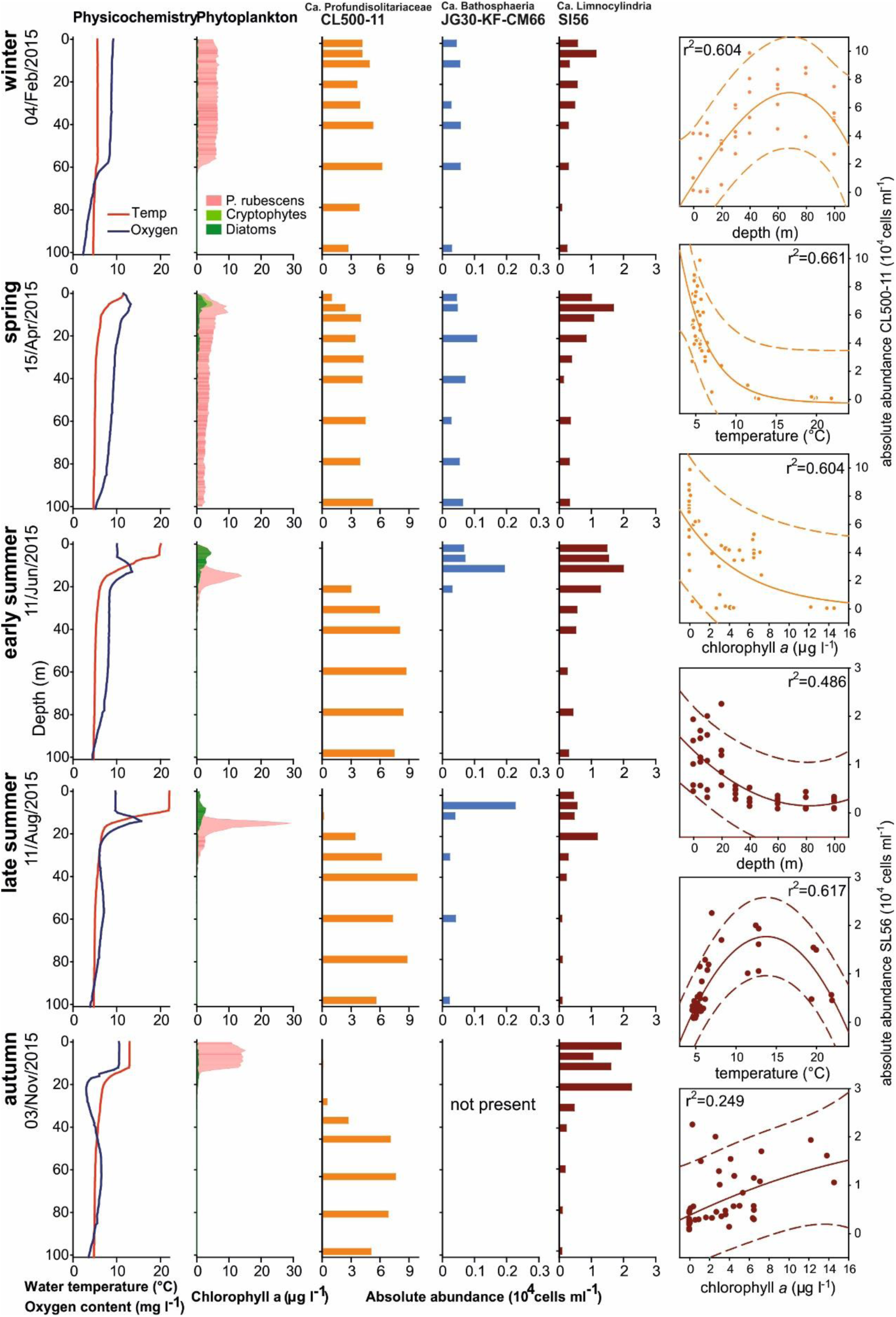
Vertical profiles of water temperature, oxygen, phytoplankton and absolute CARD-FISH abundances of three lineages of Chloroflexi in Lake Zurich at five different sampling point in 2015. Relationships of absolute abundances of the CL500-11 and SL56 groups to depth, temperature and chlorophyll *a* are shown at the right. Correlation coefficients (r^2^) are indicated within the plots.

**Supplementary Figure S4:**
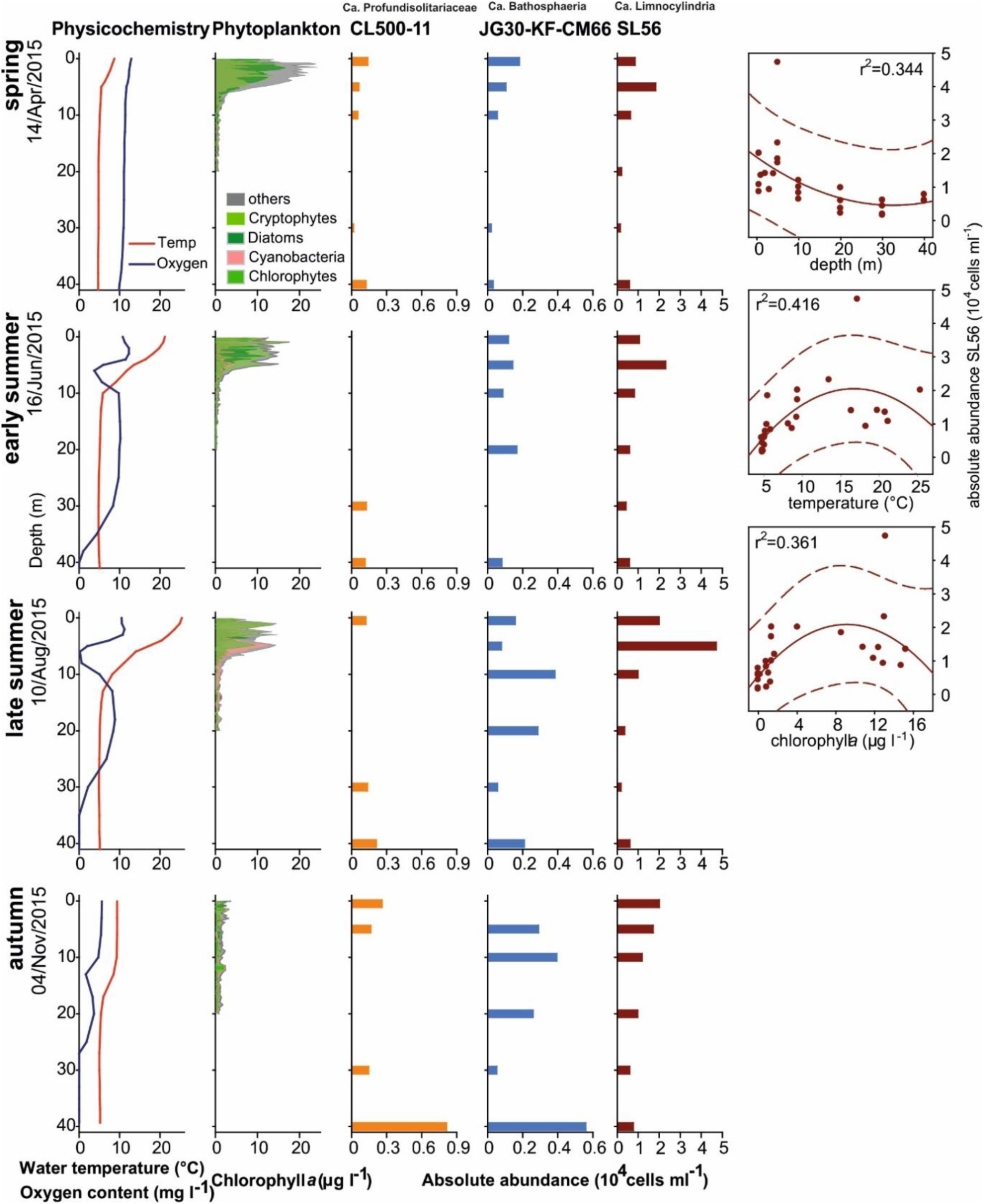
Vertical profiles of water temperature, oxygen, phytoplankton and absolute CARD-FISH abundances of three lineages of Chloroflexi in Rimov Reservoir at four different sampling points in 2015. Relationships of absolute abundance of the SL56 group to depth, temperature and chlorophyll *a* are shown at the right. Correlation coefficients (r^2^) are indicated within the plots.

**Supplementary Figure S5:**
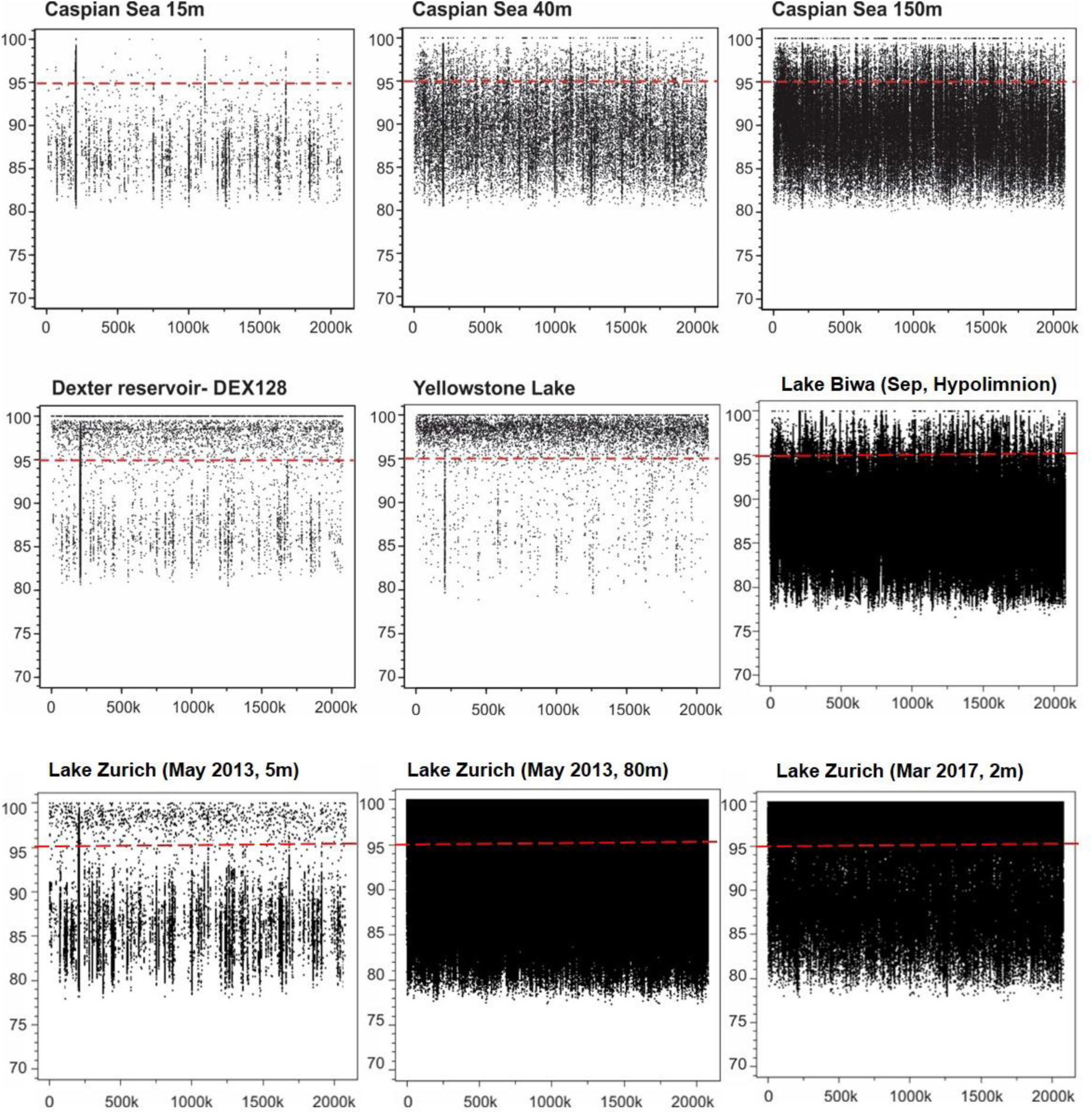
Recruitment plot for ZSMay80m-G1 as a representative of the *Chloroflexi* CL500-11 cluster against different freshwater environments and the depth profile of brackish Caspian Sea. The ZSMay80m-G1 is the only bin that contains a 16S rRNA sequence and shows completeness of 75%. In each panel the Y axis represents the identity percentage and X axis represents the genome length. The red dashed line shows the threshold for presence of same species (95% identity).

**Supplementary Figure S6:**
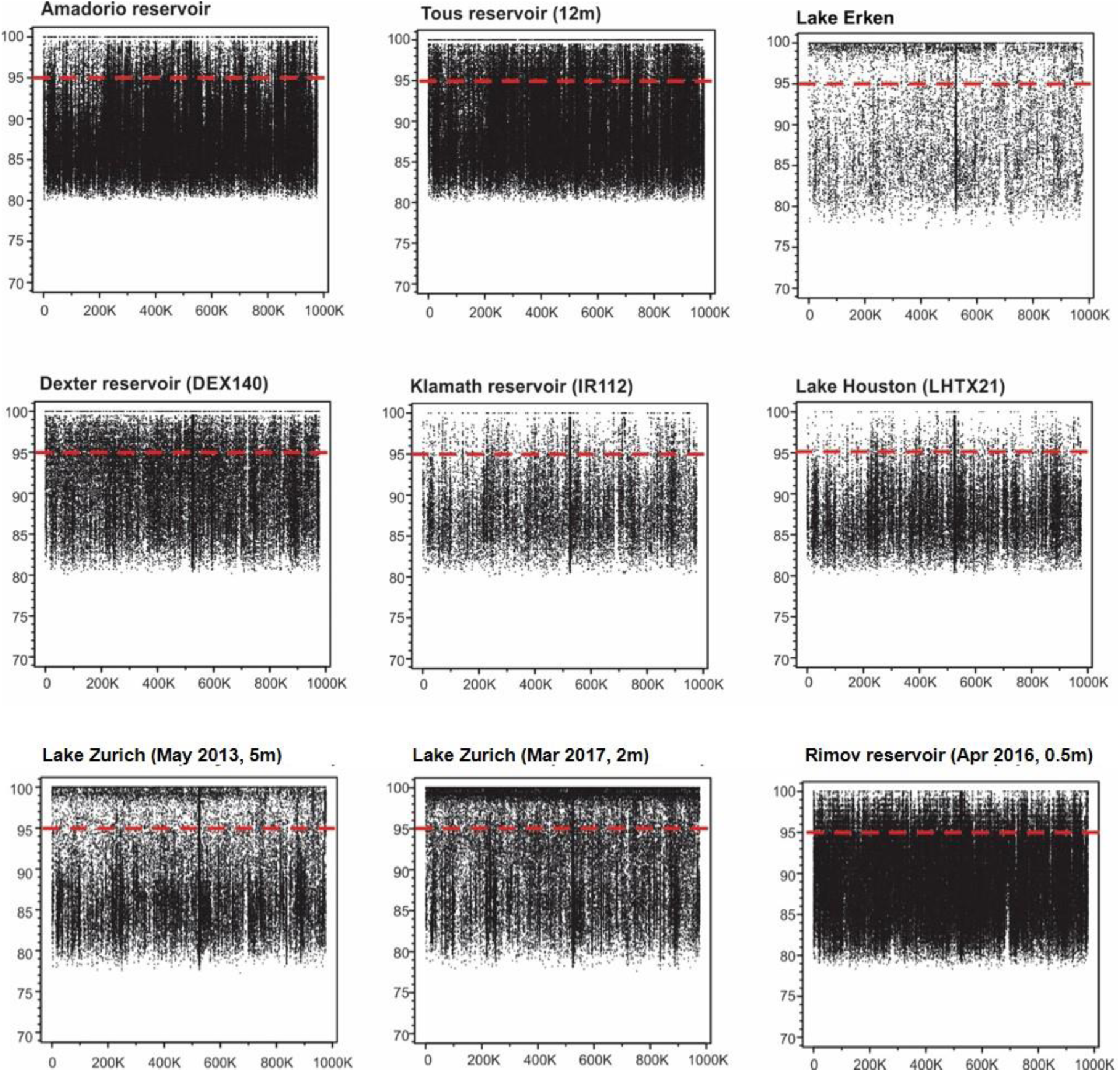
Recruitment plot for ZSMar2m-G89 as a representative of the *Chloroflexi* SL56 cluster against different freshwater environments. The ZSMar2m-G89 is the only bin that contains a 16S rRNA sequence and shows completeness of 68%. In each panel the Y axis represents the identity percentage and X axis represents the genome length. The red dashed line shows the threshold for presence of same species (95% identity).

**Supplementary Figure S7:**
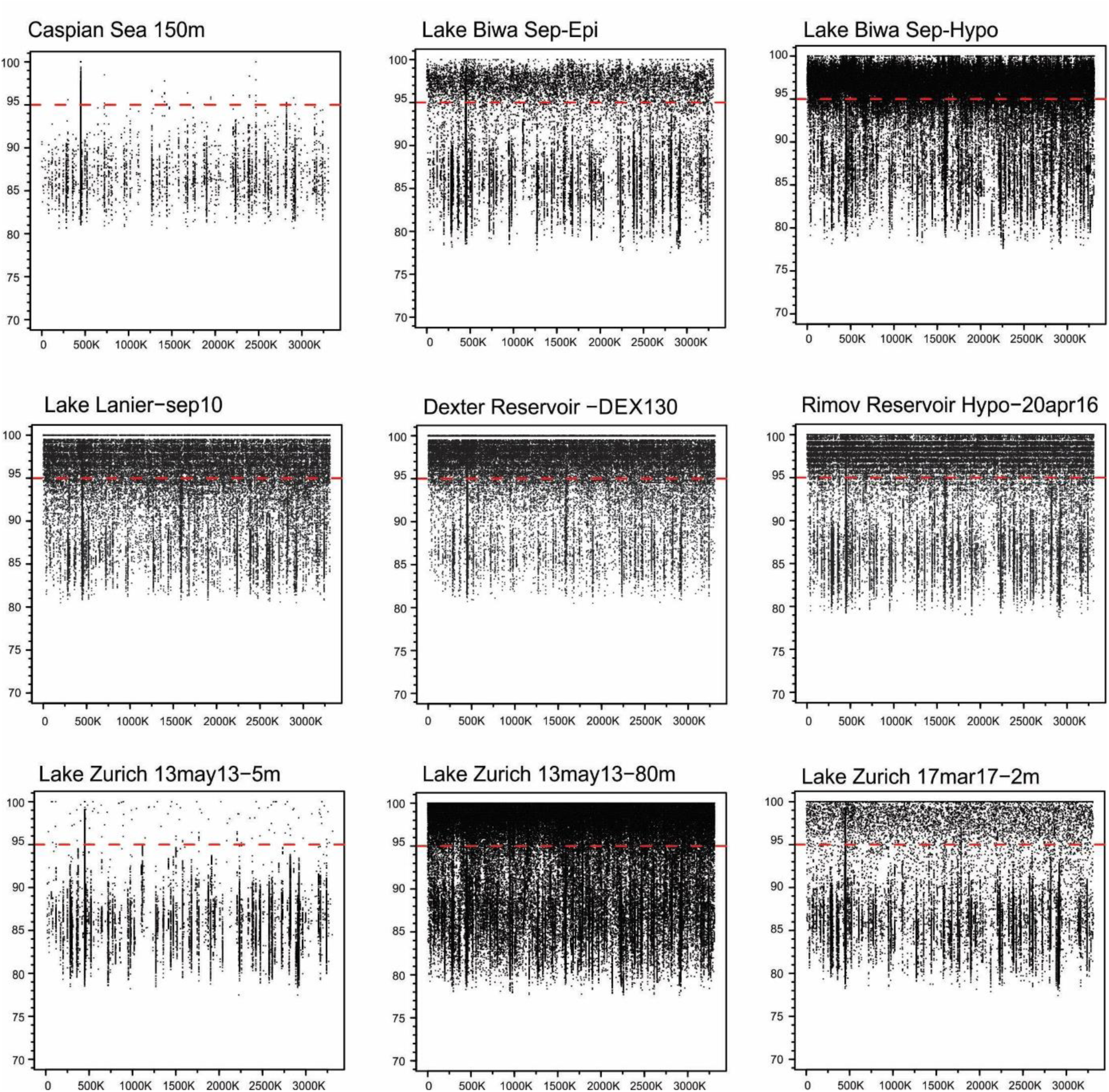
Recruitment plot for ZSMay80m-G79 as a representative of the Chloroflexi TK10 cluster against deep Caspian Sea dataset and different freshwater environments. The ZSMay80m-G79 is the most complete genome in the TK10 cluster (85%) and also contains a 16S rRNA sequence. In each panel the Y axis represents the identity percentage and X axis represents the genome length. The red dashed line shows the threshold for presence of same species (95% identity).

**Supplementary Figure S8:**
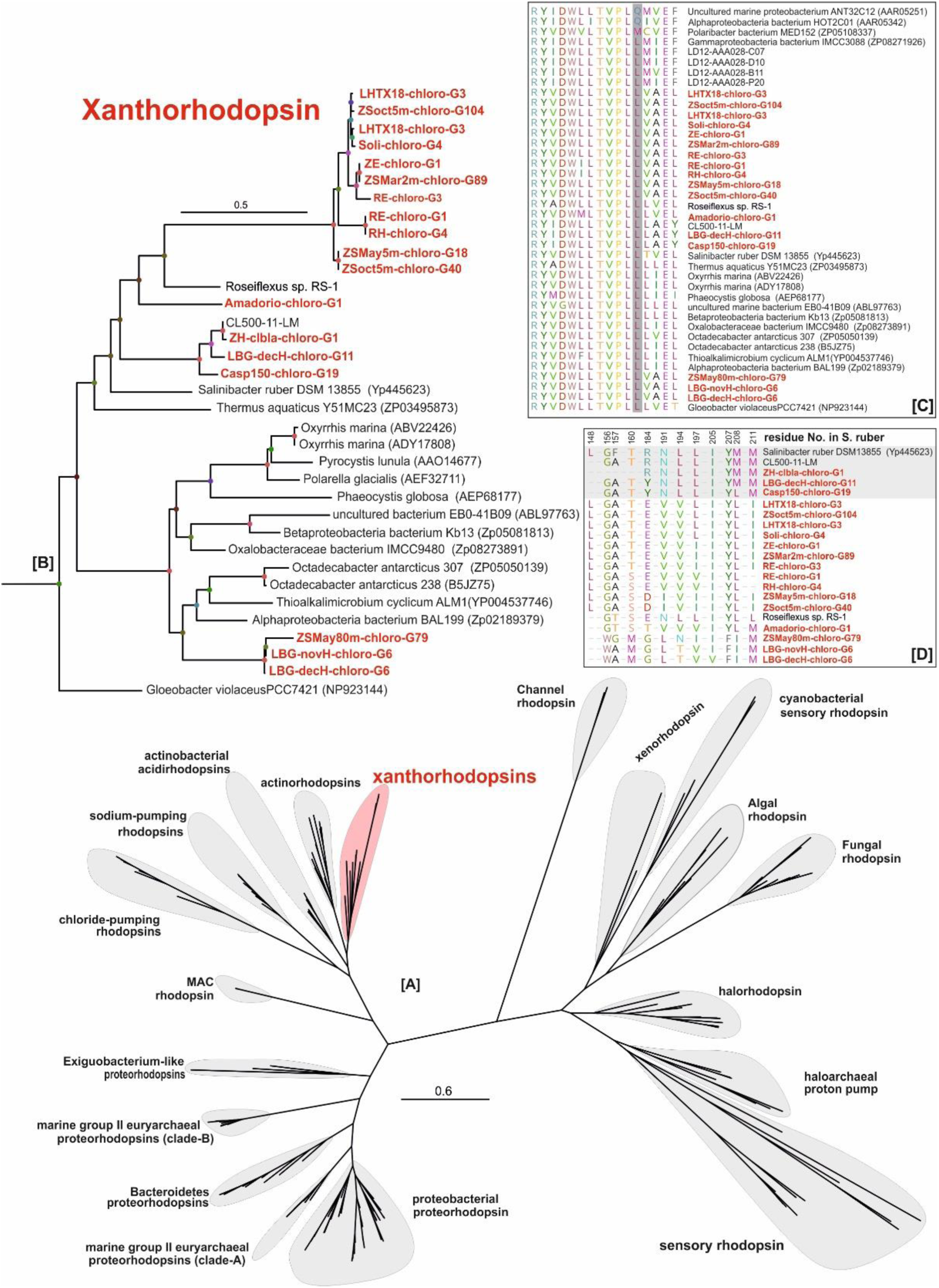
Maximum likelihood tree of rhodopsin protein sequences from different bacterial and archaeal groups (212 protein sequences in total) [A]. Expanded Maximum likelihood tree of the rhodopsin protein sequences belonging to the phylum *Chloroflexi* [B]. The alignment of the rhodopsin protein sequences from the amino acid associated with light absorption preferences. The leucine (L) and methionine (M) variants absorb maximally in the green spectrum while the glutamine (Q) variant absorbs maximally in the blue spectrum [C]. The alignment of amino acid residues involved in carotenoid binding in Salinibacter ruber DSM13855 (Luecke *et al*., 2008) and Xanthorhodopsin like sequences of the phylum *Chloroflexi*. The residue number is mentioned on top of the panel [D]. The rhodopsin genes present in the MAGs of this study are highlighted in red.

**Supplementary Figure S9:**
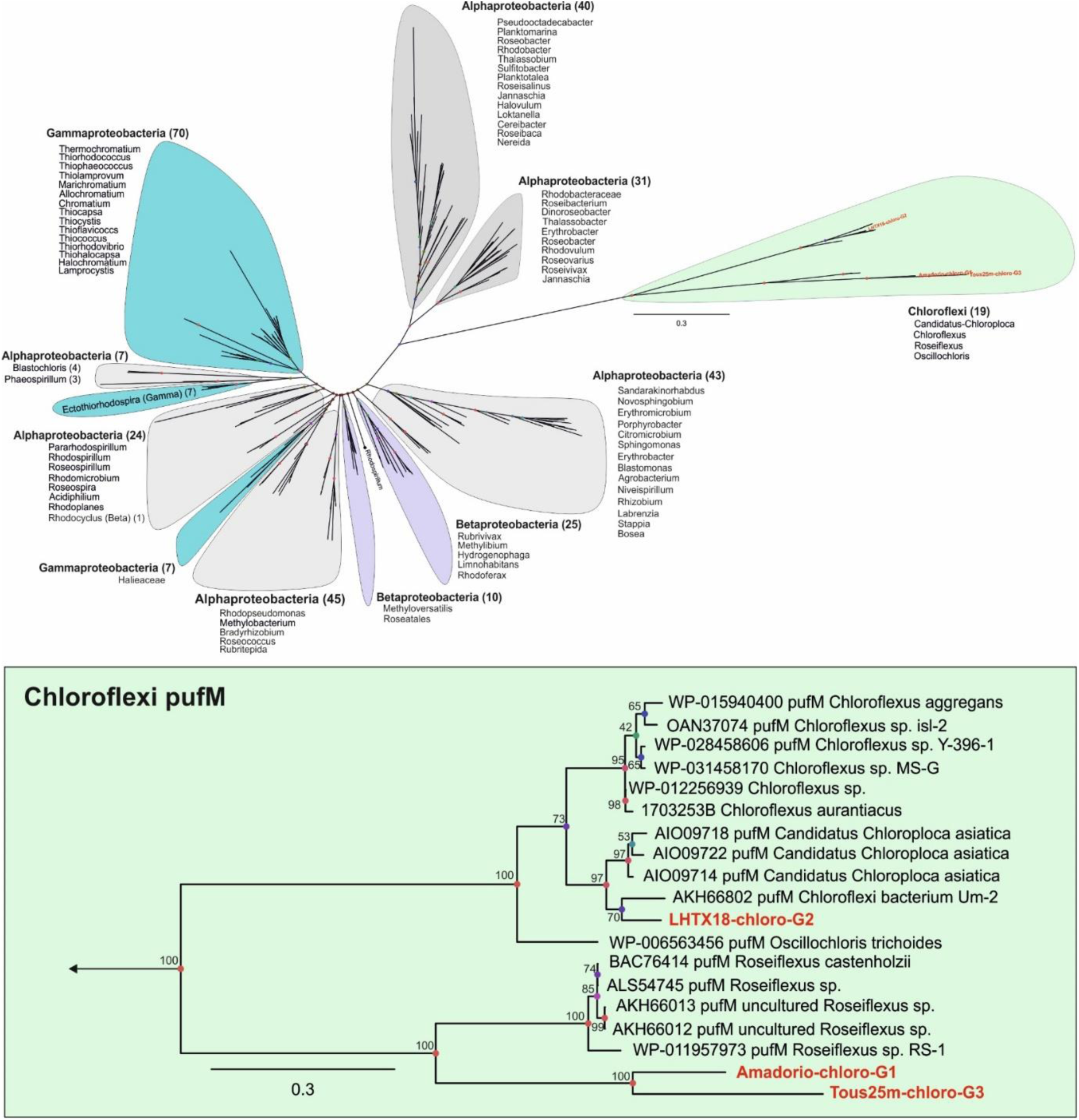
Maximum likelihood tree of the *pufM* protein sequences from different bacterial groups (328 protein sequences in total) [A]. Expanded Maximum likelihood tree of the *pufM* protein sequences belonging to the phylum *Chloroflexi* [B]. The *pufM* genes present in the MAGs of this study are highlighted in red.

## References

1. Ghai R, Mizuno CM, Picazo A, Camacho A, Rodriguez-Valera F. Key roles for freshwater Actinobacteria revealed by deep metagenomic sequencing. Mol Ecol. 2014;23:6073–90.

2. Neuenschwander SM, Ghai R, Pernthaler J, Salcher MM. Microdiversification in genome-streamlined ubiquitous freshwater Actinobacteria. Isme J. 2017;:1–14. doi:10.1038/ismej.2017.156.

3. Salcher MM, Posch T, Pernthaler J. In situ substrate preferences of abundant bacterioplankton populations in a prealpine freshwater lake. ISME J. 2013;7:896–907. doi:10.1038/ismej.2012.162.

4. Salcher MM, Neuenschwander SM, Posch T, Pernthaler J. The ecology of pelagic freshwater methylotrophs assessed by a high-resolution monitoring and isolation campaign. ISME J. 2015;9:2442–53. doi:10.1038/ismej.2015.55.

5. Kasalický V, Jezbera J, Hahn MW, Šimek K. The diversity of the Limnohabitans genus, an important group of freshwater bacterioplankton, by characterization of 35 isolated strains. PLoS One. 2013;8:e58209. doi:10.1371/journal.pone.0058209.

6. Hoetzinger M, Schmidt J, Jezberová J, Koll U, Hahn MW. Microdiversification of a pelagic Polynucleobacter species is mainly driven by acquisition of genomic islands from a partially interspecific gene pool. Appl Environ Microbiol. 2017;83:e02266–16. doi:10.1128/AEM.02266-16.

7. Salcher MM, Pernthaler J, Posch T. Seasonal bloom dynamics and ecophysiology of the freshwater sister clade of SAR11 bacteria “that rule the waves” (LD12). ISME J. 2011;5:1242–52. doi:10.1038/ismej.2011.8.

8. Henson MW, Lanclos VC, Faircloth BC, Thrash JC. Cultivation and genomics of the first freshwater SAR11 (LD12) isolate. http://dx.doi.org/101101/093567. 2016; January:1–24.

9. Ghylin TW, Garcia SL, Moya F, Oyserman BO, Schwientek P, Forest KT, et al. Comparative single-cell genomics reveals potential ecological niches for the freshwater acI Actinobacteria lineage. ISME J. 2014;8:2503–16. doi:10.1038/ismej.2014.135.

10. Cabello-Yeves PJ, Ghai R, Mehrshad M, Picazo A, Camacho A, Rodriguez-valera F. Reconstruction of diverse verrucomicrobial genomes from metagenome datasets of freshwater reservoirs. Front Microbiol. 2017.

11. Urbach E, Vergin KL, Young L, Morse A, Larson GL, Giovannoni SJ. Unusual bacterioplankton community structure in ultra-oligotrophic Crater Lake. Limnol Oceanogr. 2001;46:557–72.

12. Urbach E, Vergin KL, Larson GL, Giovannoni SJ. Bacterioplankton communities of Crater Lake, OR: Dynamic changes with euphotic zone food web structure and stable deep water populations. Hydrobiologia. 2007;574:161–77.

13. Okazaki Y, Hodoki Y, Nakano SI. Seasonal dominance of CL500-11 bacterioplankton (phylum Chloroflexi) in the oxygenated hypolimnion of Lake Biwa, Japan. FEMS Microbiol Ecol. 2013;83:82–92.

14. Okazaki Y, Nakano SI. Vertical partitioning of freshwater bacterioplankton community in a deep mesotrophic lake with a fully oxygenated hypolimnion (Lake Biwa, Japan). Environ Microbiol Rep. 2016;8:780–8.

15. Okazaki Y, Fujinaga S, Tanaka A, Kohzu A, Oyagi H. Ubiquity and quantitative significance of bacterioplankton lineages inhabiting the oxygenated hypolimnion of deep freshwater lakes. Nat Publ Gr. 2017;:1–15. doi:10.1038/ismej.2017.89.

16. Denef VJ, Mueller RS, Chiang E, Liebig JR, Vanderploeg HA. Chloroflexi CL500-11 Populations That Predominate Deep-Lake Hypolimnion Bacterioplankton Rely on Nitrogen-Rich Dissolved Organic Matter Metabolism and C ^1^ Compound Oxidation. Appl Environ Microbiol. 2016;82:1423–32. doi:10.1128/AEM.03014-15.

17. Landry Z, Swan BK, Herndl GJ, Stepanauskas R, Giovannoni SJ. SAR202 Genomes from the Dark Ocean Predict Pathways for the Oxidation of Recalcitrant Dissolved Organic Matter. MBio. 2017;8:e00413–17. doi:10.1128/mBio.00413-17.

18. Denef VJ, Fujimoto M, Berry MA, Schmidt ML. Seasonal Succession Leads to Habitat-Dependent Differentiation in Ribosomal RNA: DNA Ratios among Freshwater Lake Bacteria. Front Microbiol. 2016;7 April:1–13.

19. Tang X, Chao J, Gong Y, Wang Y, Wilhelm SW, Gao G. Spatiotemporal dynamics of bacterial community composition in large shallow eutrophic Lake Taihu: High overlap between free-living and particle-attached assemblages. Limnol Oceanogr. 2017;62:1366–82.

20. Han M, Gong Y, Zhou C, Zhang J, Wang Z, Ning K. Comparison and Interpretation of Taxonomical Structure of Bacterial Communities in Two Types of Lakes on Yun-Gui plateau of China. Nat Publ Gr. 2016; April:1–12. doi:10.1038/srep30616.

21. Pruesse E, Quast C, Knittel K, Fuchs BM, Ludwig W, Peplies J, et al. SILVA: A comprehensive online resource for quality checked and aligned ribosomal RNA sequence data compatible with ARB. Nucleic Acids Res. 2007;35:7188–96.

22. Gernert C, Glockner FO, Krohne G, Hentschel U. Microbial Diversity of the Freshwater Sponge Spongilla lacustris. Microb Ecol. 2005;50:206–12.

23. Rusch DB, Halpern AL, Sutton G, Heidelberg KB, Williamson S, Yooseph S, et al. The Sorcerer II Global Ocean Sampling expedition: northwest Atlantic through eastern tropical Pacific. PLoS Biol. 2007;5:e77. doi:10.1371/journal.pbio.0050077.

24. Morris RM, Rappé MS, Urbach E, Connon SA, Rappe MS, Giovannoni SJ. Prevalence of the Chloroflexi-Related SAR202 Bacterioplankton Cluster throughout the Mesopelagic Zone and Deep Ocean. Appl Env Microbiol. 2004;70:2836–42.

25. Schattenhofer M, Fuchs BM, Amann R, Zubkov M V., Tarran GA, Pernthaler J. Latitudinal distribution of prokaryotic picoplankton populations in the Atlantic Ocean. Environ Microbiol. 2009;11:2078–93.

26. Mehrshad M, Rodriguez-Valera F, Amoozegar MA, López-García P, Ghai R. The enigmatic SAR202 cluster up close: shedding light on a globally distributed dark ocean lineage involved in sulfur cycling. ISME J. 2017.

27. Lozupone CA, Knight R. Global patterns in bacterial diversity. Proc Natl Acad Sci. 2007;104:11436–40.

28. Walsh DA, Lafontaine J, Grossart H-P. On the Eco-Evolutionary Relationships of Fresh and Salt Water Bacteria and the Role of Gene Transfer in Their Adaptation. In: Gophna U, editor. Lateral Gene Transfer in Evolution. New York, NY: Springer New York; 2013. p. 55–77. doi:10.1007/978-1-4614-7780-8_3.

29. Salcher MM, Neuenschwander SM, Posch T, Pernthaler J. The ecology of pelagic freshwater methylotrophs assessed by a high-resolution monitoring and isolation campaign. ISME J. 2015;9:2442–53. doi:10.1038/ismej.2015.55.

30. Logares R, Bråte J, Bertilsson S, Clasen JL, Shalchian-Tabrizi K, Rengefors K. Infrequent marine-freshwater transitions in the microbial world. Trends Microbiol. 2009;17:414–22. doi:http://dx.doi.org/10.1016/j.tim.2009.05.010.

31. Eiler A, Mondav R, Sinclair L, Fernandez-Vidal L, Scofield DG, Schwientek P, et al. Tuning fresh: Radiation through rewiring of central metabolism in streamlined bacteria. ISME J. 2016;10:1902–14. doi:10.1038/ismej.2015.260.

32. Posch T, Köster O, Salcher MM, Pernthaler J. Harmful filamentous cyanobacteria favoured by reduced water turnover with lake warming. Nat Clim Chang. 2012;2:809–13. doi:10.1038/nclimate1581.

33. Hug LA, Thomas BC, Sharon I, Brown CT, Sharma R, Hettich RL, et al. Critical biogeochemical functions in the subsurface are associated with bacteria from new phyla and little studied lineages. Environ Microbiol. 2016;18:159–73.

34. Tripp HJ, Kitner JB, Schwalbach MS, Dacey JWH, Wilhelm LJ, Giovannoni SJ. SAR11 marine bacteria require exogenous reduced sulphur for growth. Nature. 2008;452:741–4. doi:10.1038/nature06776.

35. Doxey AC, Kurtz D a, Lynch MD, Sauder L a, Neufeld JD. Aquatic metagenomes implicate Thaumarchaeota in global cobalamin production. ISME J. 2014;:1–11. doi:10.1038/ismej.2014.142.

36. Qin W, Amin SA, Lundeen RA, Heal KR, Martens-habbena W, Turkarslan S, et al. Stress response of a marine ammonia-oxidizing archaeon informs physiological status of environmental populations. Nat Publ Gr. 2017; June:1–12. doi:10.1038/ismej.2017.186.

37. Roth JR, Lawrence JG, Bobik TA. COBALAMIN (COENZYME B 12): Synthesis and Biological Significance. 1996.

38. Morris JJ, Lenski RE, Zinser ER. The Black Queen Hypothesis: Evolution of Dependencies through Adaptive Gene Loss. MBio. 2012;3:1–7.

39. Men Y, Seth EC, Yi S, Allen RH, Taga ME, Alvarez-cohen L. Sustainable Growth of Dehalococcoides mccartyi 195 by Corrinoid Salvaging and Remodeling in Defined Lactate-Fermenting Consortia. Appl Environmantal Microbiol. 2014;80:2133–41.

40. Escalante-Semerena JC. Conversion of cobinamide into adenosylcobamide in bacteria and archaea. J Bacteriol. 2007;189:4555–60.

41. Wu D, Raymond J, Wu M, Chatterji S, Ren Q, Graham JE, et al. Complete genome sequence of the aerobic CO-oxidizing thermophile Thermomicrobium roseum. PLoS One. 2009;4:e4207.

42. Sutcliffe IC. Cell envelope architecture in the Chloroflexi: A shifting frontline in a phylogenetic turf war. Environ Microbiol. 2011;13:279–82.

43. Balashov SP, Imasheva ES, Boichenko V a, Antón J, Wang JM, Lanyi JK. Xanthorhodopsin: a proton pump with a light-harvesting carotenoid antenna. Science. 2005;309:2061–4. doi:10.1126/science.1118046.

44. Balashov SP, Lanyi JK. Xanthorhodopsin: Proton pump with a carotenoid antenna. Cell Mol Life Sci. 2007;64:2323–8.

45. Boichenko VA, Wang JM, Antón J, Lanyi JK, Balashov SP. Functions of Carotenoids in Xanthorhodopsin and Archaerhodopsin, from Action Spectra of Photoinhibition of Cell Respiration. Biochim Biophys Acta. 2006;1757:1649–1656.

46. Zeng Y, Feng F, Medová H, Dean J, Koblízek M. Functional type 2 photosynthetic reaction centers found in the rare bacterial phylum Gemmatimonadetes. Pnas. 2014;111:7795–800.

47. Walsh DA, Lafontaine J, Grossart H-P. On the Eco-Evolutionary Relationships of Fresh and Salt Water Bacteria and the Role of Gene Transfer in Their Adaptation. In: Gophna U, editor. Lateral Gene Transfer in Evolution. New York, NY: Springer New York; 2013. p. 55–77.

48. Simek K, Bobková J, Macek M, Nedoma J, Psenner R. Ciliate grazing on picoplankton in a eutrophic reservoir during the summer phytoplankton maximum: A study at the species and community level. Limnol Oceanogr. 1995;40:1077–90.

49. Martín-Cuadrado A-B, López-García P, Alba J-C, Moreira D, Monticelli L, Strittmatter A, et al. Metagenomics of the deep Mediterranean, a warm bathypelagic habitat. PLoS One. 2007;2:e914. doi:10.1371/journal.pone.0000914.

50. Edgar RC. Search and clustering orders of magnitude faster than BLAST. Bioinformatics. 2010;26:2460–1. doi:10.1093/bioinformatics/btq461.

51. Nawrocki E. Structural RNA Homology Search and Alignment Using Covariance Models. Washington University in ST. Louis; 2009.

52. Altschul SF, Madden TL, Schäffer AA, Zhang J, Zhang Z, Miller W, et al. Gapped BLAST and PSI-BLAST: a new generation of protein database search programs. Nucleic Acids Res. 1997;25:3389–402.

53. Pruesse E, Peplies J, Glöckner FO. SINA: Accurate high-throughput multiple sequence alignment of ribosomal RNA genes. Bioinformatics. 2012;28:1823–9.

54. Ludwig W, Strunk O, Westram R, Richter L, Meier H, Yadhukumar A, et al. ARB: A software environment for sequence data. Nucleic Acids Res. 2004;32:1363–71.

55. Stamatakis A, Ludwig T, Meier H. RAxML-II: A program for sequential, parallel and distributed inference of large phylogenetic trees. Concurr Comput Pract Exp. 2005;17:1705–23.

56. Zeder M, Pernthaler J. Multispot live-image autofocusing for high-throughput microscopy of fluorescently stained bacteria. Cytom Part A. 2009;75:781–8.

57. Posch T, Franzoi J, Prader M, Salcher MM. New image analysis tool to study biomass and morphotypes of three major bacterioplankton groups in an alpine lake. Aquat Microb Ecol. 2009;54:113–26.

58. Nurk S, Meleshko D, Korobeynikov A, Pevzner PA. metaSPAdes: a new versatile metagenomic assembler. 2017;:824–34.

59. Bolger AM, Lohse M, Usadel B. Genome analysis Trimmomatic: a flexible trimmer for Illumina sequence data. 2014;30:2114–20.

60. Schmieder R, Edwards R. Quality control and preprocessing of metagenomic datasets. Bioinformatics. 2011;27:863–4.

61. Hyatt D, Chen G-L, LoCascio PF, Land ML, Larimer FW, Hauser LJ. Prodigal: prokaryotic gene recognition and translation initiation site identification. BMC Bioinformatics. 2010;11:119. doi:10.1186/1471-2105-11-119.

62. Steinegger M, Söding J. MMseqs2 enables sensitive protein sequence searching for the analysis of massive data sets. Nat Biotechnol. 2017;35:1026–1028.

63. Kang DD, Froula J, Egan R, Wang Z. MetaBAT, an efficient tool for accurately reconstructing single genomes from complex microbial communities. PeerJ. 2015;3:e1165. doi:10.7717/peerj.1165.

64. Seemann T. Prokka: Rapid prokaryotic genome annotation. Bioinformatics. 2014;30:2068–9.

65. Tatusov RL, Natale DA, Garkavtsev I V, Tatusova TA, Shankavaram UT, Rao BS, et al. The COG database: new developments in phylogenetic classification of proteins from complete genomes. Nucleic Acids Res. 2001;29:22–8.

66. Haft DH, Loftus BJ, Richardson DL, Yang F, Eisen JA, Paulsen IT, et al. TIGRFAMs: a protein family resource for the functional identification of proteins. Nucleic Acids Res. 2001;29:41–3.

67. Eddy SR. Accelerated profile HMM searches. PLoS Comput Biol. 2011;7:e1002195.

68. Aziz RK, Bartels D, Best A a, DeJongh M, Disz T, Edwards R a, et al. The RAST Server: rapid annotations using subsystems technology. BMC Genomics. 2008;9:75. doi:10.1186/1471-2164-9-75.

69. Kanehisa M, Sato Y, Morishima K. BlastKOALA and GhostKOALA: KEGG Tools for Functional Characterization of Genome and Metagenome Sequences. J Mol Biol. 2016;428:726–31. doi:10.1016/j.jmb.2015.11.006.

70. Claudel-Renard C, Chevalet C, Faraut T, Kahn D. Enzyme-specific profiles for genome annotation: PRIAM. Nucleic Acids Res. 2003;31:6633–9.

71. Parks DH, Imelfort M, Skennerton CT, Hugenholtz P, Tyson GW. CheckM: assessing the quality of microbial genomes recovered from isolates, single cells, and metagenomes. Genome Res. 2015;25:1043–55. doi:10.1101/gr.186072.114.

72. Segata N, Börnigen D, Morgan XC, Huttenhower C. PhyloPhlAn is a new method for improved phylogenetic and taxonomic placement of microbes. Nat Commun. 2013;4:2304. doi:10.1038/ncomms3304.

73. Edgar RC. MUSCLE: multiple sequence alignment with high accuracy and high throughput. Nucleic Acid Res. 2004;32:1792–7.

74. Price MN, Dehal PS, Arkin AP. FastTree 2--approximately maximum-likelihood trees for large alignments. PLoS One. 2010;5:e9490. doi:10.1371/journal.pone.0009490.

75. Shimodaira H, Hasegawa M. Multiple Comparisons of Log-Likelihoods with Applications to Phylogenetic Inference. Mol Biol Evol. 1999;16:1114–6.

76. Konstantinidis KT, Tiedje JM. Genomic insights that advance the species definition for prokaryotes. Proc Natl Acad Sci U S A. 2005;102:2567–72. doi:10.1073/pnas.0409727102.

